# Nanoscale Structural and Functional Impacts of Disease-Associated Collagen Mutations

**DOI:** 10.1101/2025.09.03.673992

**Authors:** Caitlyn A. Tobita, Siddhartha Banerjee, Jonathan Roth, Emma K. Larson, Paulina R. Fernandez, Abuzar Nikzad, Abdullah Naiyer, Cody L. Hoop, Jean Baum

## Abstract

Collagen is the most abundant structural protein in the human body, and its supramolecular organization is central to tissue mechanics and cell–matrix interactions. Integrins, key mediators of these interactions, are essential for important biological processes including adhesion, migration, differentiation, and platelet aggregation. While mutations in collagen are known to cause connective tissue disorders such as Osteogenesis Imperfecta (OI) with phenotypes ranging from mild to perinatal lethal, how these mutations alter fibril level architecture, dynamics, integrin-mediated interactions and their functional consequences at both the protein and cellular level remains poorly understood. Here, we investigated the impact of OI mutations on direct collagen-integrin interactions and on cellular responses to mutant collagen. We generated collagen-rich extra-cellular matrix (ECM) from primary dermal fibroblasts of a healthy donor (WT) and from two OI patients carrying distinct glycine mutations: G610C, associated with moderate disease, and G907D, linked to perinatal lethality. Comparative biophysical studies reveal that both mutants retain the canonical D-banding of collagen I fibrils but differ markedly at the nanoscale. G907D fibrils exhibit greater local structural perturbations and increased molecular mobility relative to the non-lethal G610C. Importantly, while cell surface adhesion and proliferation do not differ between the OI mutants, integrin binding diverges between mutants: G610C displays reduced affinity, whereas G907D exhibits enhanced affinity compared to WT. Together, these findings establish a mechanistic link between single-residue mutations, nanoscale fibril architecture and collagen-receptor interactions, and highlight how genetic or acquired collagen defects can drive ECM dysregulation.

## Introduction

The extracellular matrix (ECM) is a dynamic and essential component of multicellular life, providing not only structural support to tissues but also regulating critical biological processes such as cell adhesion, migration, differentiation, and response to injury^1-3^. Central to the ECM’s mechanical and signaling functions are fibrillar collagens, which assemble into complex hierarchical structures that maintain tissue integrity and coordinate cellular behavior^4^. Disruptions to collagen structure and assembly underlie a wide array of connective tissue diseases, including osteogenesis imperfecta (OI), Ehlers-Danlos syndrome (EDS), and Marfan syndrome^5-7^. These conditions can lead to profound consequences ranging from skeletal fragility to cardiovascular complications, affecting individuals across all age groups.

Osteogenesis imperfecta (OI), also known as brittle bone disease, is a heritable connective tissue disorder characterized by fragile bones, frequent fractures, and, in severe cases, perinatal lethality. The majority of OI cases result from dominant-negative mutations in *COL1A1* or *COL1A2* genes, which encode type I collagen, the most abundant structural protein in the human body. OI affects an estimated 50,000 people in the United States alone and is clinically classified into different types based on symptom severity^9^. Most dominant-negative OI mutations involve substitutions of glycine residues within the repeating Gly-X-Y (GXY, X/Y represents variable amino acid) motif that is essential for the stability of the collagen triple helix. In this study, we focus on two clinically distinct COL1A2 glycine substitutions, G610C, associated with moderate disease severity, and G907D, associated with a perinatal lethal phenotype. The severity of the disease often depends on the identity and position of the substituted residue, including the size, charge and local sequence context^8-12^, but the molecular mechanisms that connect these mutations to altered collagen structure and downstream cell-matrix signaling remain poorly understood.

Type I collagen assembles into highly ordered fibrils with a characteristic 67 nm D-banding pattern formed by staggered packing of triple-helical monomers. The repeating GXY amino acid sequence of collagen assembles into a three-chain triple helix monomer (Fig. 1A). These triple-helical monomers self-assemble in a uniform, staggered arrangement to form fibrils (Fig. 1B). The staggered alignment of five monomers creates distinct overlap and gap regions, which repeat in three dimensions and give rise to the periodic D-banding (Fig.1B). While atomic-resolution studies using collagen-mimetic peptides have revealed that glycine substitutions introduce local kinks and dynamic perturbations into the triple helix^13-16^, it is not yet clear how these local disruptions propagate to affect the nanoscale architecture and functional interactions of full-length collagen fibrils in a physiologically relevant context. This represents a critical gap, as the spatial arrangement and dynamic behavior of collagen fibrils directly influence their interactions with other ECM components and cell surface receptors.

**Figure 1:**
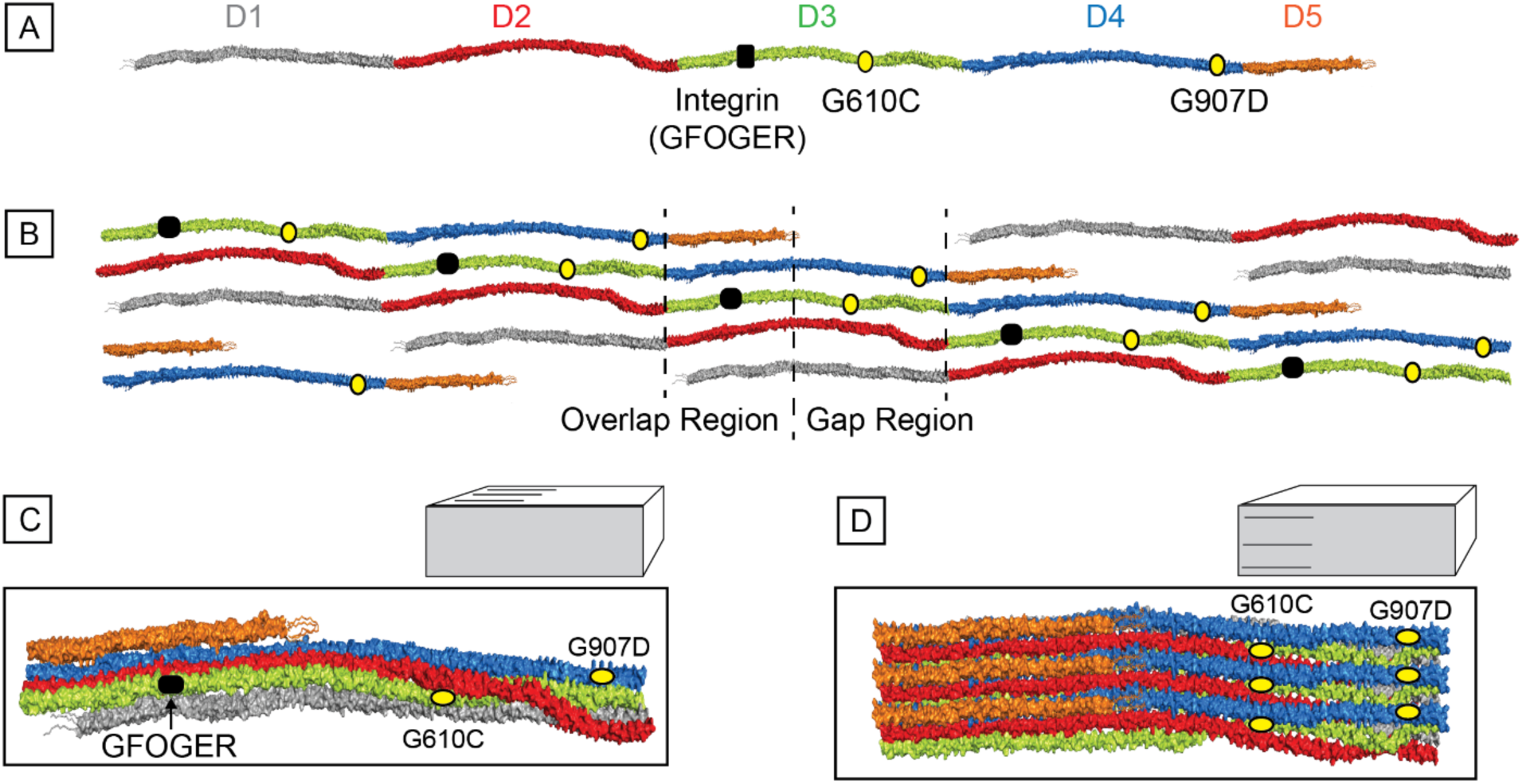
Collagen I assembly, integrin binding region, and the location of OI mutations in the collagen monomer. (A) The collagen monomers are composed of 3 polypeptide chains, two a1 chains and one a2 chain for collagen I, that are supercoiled to form a triple helix (surface view). The monomer is divided into 5 D-periods, which are colored as follows: D1(grey), D2 (red), D3 (green), D4 (blue), and D5 (orange). The black box marks the location of the high affinity integrin I domain binding site, GFOGER^71^. OI mutations G610C (non-lethal)and G907D (lethal), located in D3 and D4 respectively, are indicated by yellow circles. (B) Schematic of collagen fibril assembly. Collagen monomers assemble in a staggered, axial arrangement such that each monomer is offset from its neighbor by one D-period. This staggered packing creates characteristic regions along the fibril known as the gap and overlap regions which correspond to areas with lower and higher monomer density respectively. A single repeating unit and the corresponding gap and overlap regions are outlined with dashed lines (C-D) Collagen fibrils form three-dimensional structures, which wind together, forming a more complex structure compared to the simplified cartoon in B. (C) Side view of the fibril, showcasing the location of the integrin binding site relative to the sites of the mutations studied. (D) Surface views of the model with three units adjacent to one another. The GFOGER site is not visible in this surface representation as it is buried beneath triple helices and thus is not shown. All structures shown are based off a molecular dynamics simulations previously done by our lab^22,72^.

One such receptor, the α2β1 integrin, plays a pivotal role in collagen-mediated cell adhesion and signaling and is a primary receptor for type I collagen^17,18^. The GXY motif known to be the integrin-binding sites, are distributed throughout the collagen α-chains^19^. Among these, the GFOGER sequence has been identified as having particularly strong binding affinity to α2β1 integrin^20^. Fig. 1 illustrates the location of GFOGER on the type I collagen triple helical monomer (Fig. 1A/B) and the subsequent assembly of monomers into the fibril structure (Fig. 1C/D). During fibril assembly, many integrin-binding sites become buried, limiting their accessibility (Fig 1C-D). Our lab has previously investigated the dynamic nature of the fibrillogenesis process as well as the dynamics of fibrils, which exhibited possibility that, through surface dynamics, these cryptic binding sites can become accessible^21-23^. However, collagen in diseased states may perturb these dynamics or the structure of the fibril surface and thus disrupt how collagen fibrils regulate activity with binding partners^24,25^.

Although OI has been studied across a range of biological scales, from peptide models to mouse systems, direct nanoscale investigation of human-derived mutant collagen fibrils remains scarce. At the molecular level, collagen mimetic peptides have been used to establish how OI mutations disrupt processes such as triple helix folding^26-28^ or internal hydrogen bonding^29,30^. For nanoscale studies, bones and tendons obtained from OI mouse models are used to investigate the effects of mutations on the fibril mineralization and mechanical properties,^31-33^ and on the microscale, several studies have performed electron microscopy to determine how the bones of OI patients are affected^25,34-36^. Yet, there remains a critical gap: there is a distinct lack of nanoscale collagen fibril studies using collagens derived from human OI patients in non-clinical, *in vitro* settings. Although studies using collagen mimetic peptides provided valuable insights into structural alterations of the collagen triple helix caused by OI mutations^13-15,37^, they lack the native structural and mechanical complexity of the extracellular matrix environment of collagen. As a result, peptide studies do not offer insights into collagen’s natural fibrillar organization or its dynamic interaction with integrin and other ECM components.

In this study, we address this challenge by analyzing type I collagen fibrils produced from primary dermal fibroblasts of healthy donors and individuals with OI. We focus on two clinically distinct glycine substitutions: G610C, associated with a moderate disease severity, and G907D, linked to perinatal lethal phenotype. Using a multimodal approach that includes transmission electron microscopy (TEM), atomic force microscopy imaging (AFM), solid state NMR (ssNMR), integrin-binding assays, and cell-based adhesion and proliferation assays, we compare the structure, dynamics and cell interaction potential of wild-type and mutant collagen fibrils from the molecular to the nanoscale level. We show that the G907D mutation induces pronounced nanoscale alterations in fibril morphology and dynamics, leading to enhanced exposure to integrin binding sites while G610C retains WT-like surface topography but very limited integrin binding ability. Despite these mutation dependent differences in collagen-integrin interaction, cell adhesion and proliferation on mutant collagen ECM remain comparable to WT. Together, these results demonstrate how specific glycine substitutions propagate structural changes along the collagen triple helix to reshape fibril surface dynamics and modulate receptor binding. Importantly, they establish a direct molecular link between OI-associated collagen mutations, fibril architecture, and integrin accessibility, providing a new mechanistic insight into how ECM alterations drive connective tissue dysfunction.

## Results

### Nanoscale morphological differences in fibroblast-derived WT and OI-mutant collagen I fibrils

In this study, fibroblast cells from healthy donors and individuals with OI were cultured to produce collagen-rich extracellular matrices (ECMs) for nanoscale analysis. The resulting ECMs, containing native and OI collagen fibrils, were harvested, gently sonicated, and used in subsequent experiments. Samples from both healthy and OI-derived ECMs were prepared for atomic force microscopy (AFM) to assess fibril morphology by measuring surface topography, fibril height and D-periodicity. Wild-type (WT) fibrils displayed uniform topography with clearly defined and regular D-banding (Fig. 2A). The G610C mutant fibrils appeared largely similar to WT, exhibiting no significant changes in D-periodicity and fibril height (Fig. 2B). In contrast, fibrils from the G907D mutant exhibited pronounced local morphological abnormalities, including fibril thinning (Fig. 2C). Despite these morphological differences, the characteristic D-band periodicity observed in WT fibrils is preserved across both OI mutants, as shown in the AFM box plots (Fig. 2D). However, the height of the G907D mutant fibrils (12.0 ± 3.5 nm) are significantly reduced compared to both WT (19.5 ± 4.3 nm) and G610C (17.3 ± 4.1 nm) which showed statistically similar height distributions. Quantitative measurements of fibril height and periodicity are provided in Supplementary Table 1 and indicate that the G907D mutation causes significant structural defects leading to the generation of abnormal collagen fibrils.

**Figure 2:**
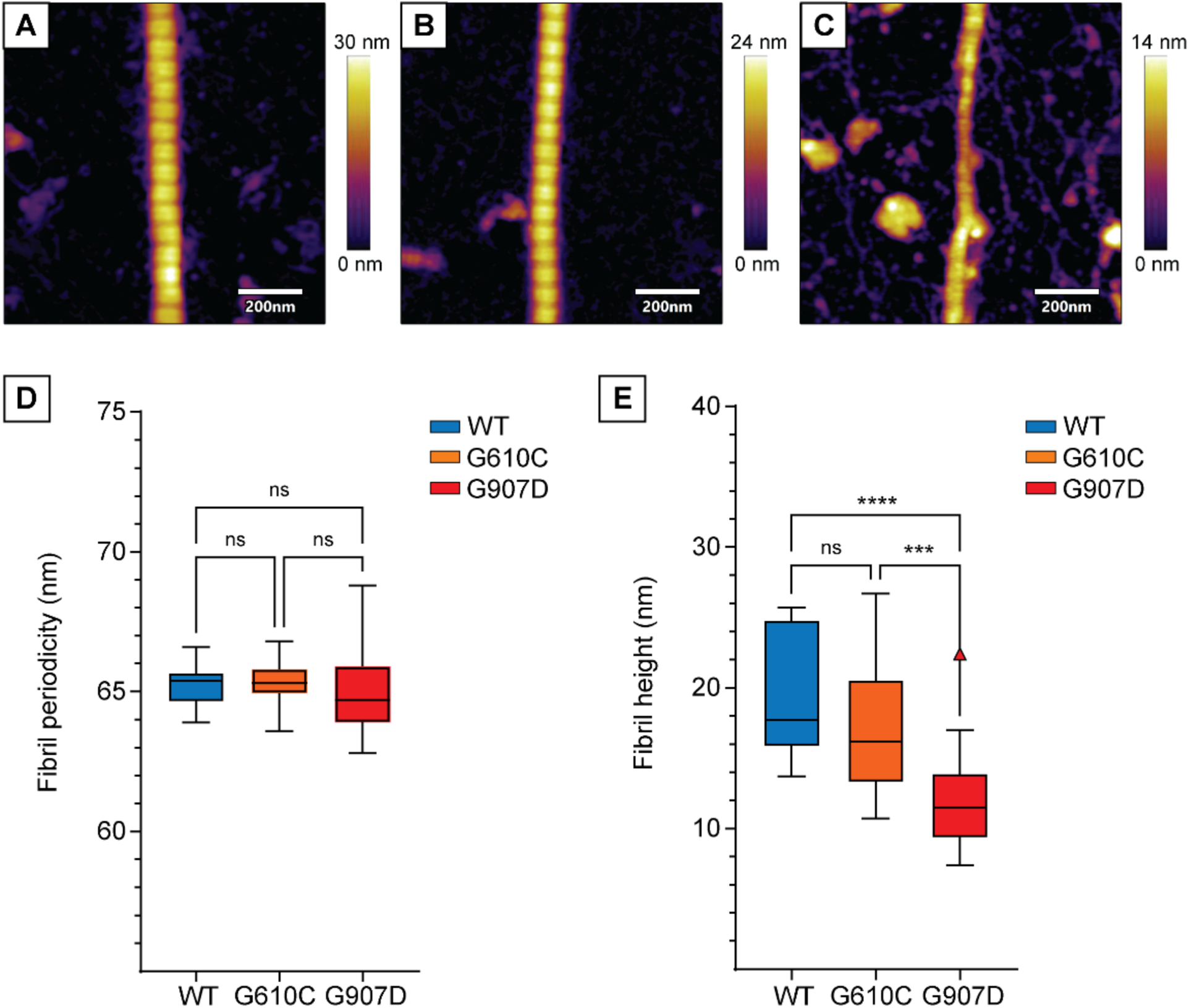
AFM analysis of human fibroblast produced collagen. Representative 1 µm x 1 µm AFM height images of (A) WT, (B) G610C mutated, and (C) G907D mutated collagen fibrils on freshly cleaved mica imaged in air. WT and G610C fibrils were extremely regular, with clear D-banding and minimal variations along fibril axis. On the other hand, G907D fibrils showed localized regions of fibril thinning compared to both WT and G610C fibrils. (D-E) Box and whisker plots of (D) fibril periodicity and (E) fibril heights, as seen in the AFM images. The statistics from D and E are available in Table S1. The box and whisker plots show the average, 25^th^, and 75^th^ percentiles in the box, and uses a Tukey method to plot for the whiskers and outliers.

To complement the surface topography obtained with AFM, which primarily provides quantitative height information, we employed TEM to visualize fibril ultrastructure and lateral morphology at high-resolution. The WT, G610C, and G907D fibrils all exhibited the characteristic D-banding pattern, reflecting the repeating overlap and gap regions of the fibril structure (Supplementary Fig. S1). WT and G610C fibrils displayed regular D-banding features, including bright white bands between each gap and overlap region, which are attributed to telopeptides and sub-bands formed by variations in charge alignment within the fibril^38^. In contrast, G907D fibrils showed local fibril thinning and reduced fibril width consistent with the altered morphology observed by AFM (Supplementary Fig. S1, S2 and Table S2). Together, the TEM and AFM data reveal that the G907D mutation introduces nanoscale structural defects not present in WT or G610C fibrils, highlighting a potential structural basis for its more severe phenotype.

### Conformational dynamics and structural alterations in collagen mutants revealed by solid-state NMR

Structural and dynamic features of the WT and mutant fibrils were compared using solid-state NMR (ssNMR). Cross polarization (CP) based experiments enhance signals of rigid components of the ECM, while Insensitive Nuclei Enhanced by Polarization Transfer (INEPT) based experiments are more sensitive to mobile components. For WT and mutant ECM production, cell cultures were enriched with [13C,15N]-Gly, Pro, and Hydroxyproline (Hyp) for detection by NMR. Gly, Pro, and Hyp in WT ECM shows characteristic chemical shift signatures for collagen I^39-41^ in the ^1^H-^13^C 1D CP spectrum (Fig. 3A, Supplementary Table S3). These chemical shifts are maintained in G610C and G907D mutant ECM (Fig. 3A, Supplementary Table S3). By 2D ^13^C-^13^C ssNMR, two conformations of Pro are apparent (Supplementary Fig. S3), which are attributed to Pro preceding a Hyp (P_GPO_) and Pro preceding any other amino acid (P_GPX_), based on assignments of analogously labeled ECM from MC3T3 cells^42^ (Supplementary Fig. S4). However, G907D ECM shows noticeably reduced signal intensities relative to WT and G610C ECM. This is complemented by the increased signal intensity for all labeled amino acids in G907D ECM in the ^13^C-INEPT 1D, suggesting that G907D ECM may be more mobile (Fig. 3B). Conversely, little to no signal is detectable in the ^13^C INEPT of WT or G610C ECM under the same conditions, suggesting that these sites are more rigid relative to G907D ECM. In addition, distinct Pro and Hyp Cα chemical shifts are observed for the rigid species and mobile species in G907D ECM, showing that the G907D mutated sample exhibits unique conformations compared to WT and non-lethal G610C ECM (Supplementary Table S3). Comparison between G907D ECM Pro Cα chemical shifts observed in CP-based and INEPT spectra reveal differences in secondary structures between the rigid and mobile Pro conformations. To contextualize the distribution of G907D ECM Pro Cα chemical shifts from rigid (red dashed lines) or mobile (blue dashed lines) Pro, we plotted them against all Pro Cα chemical shifts reported in the BMRB (Supplementary Fig. S5B) and against the subset of Pro that have PPII-like secondary structure (Supplementary Fig. S5C). We find that the rigid Pro Cα’s have chemical shifts more characteristic of those in a PPII structure, while the mobile Pro Cα’s have uniquely downfield chemical shifts. This suggests that G907D substitutions induce mobile regions within collagen I that do not maintain PPII structure.

**Figure 3:**
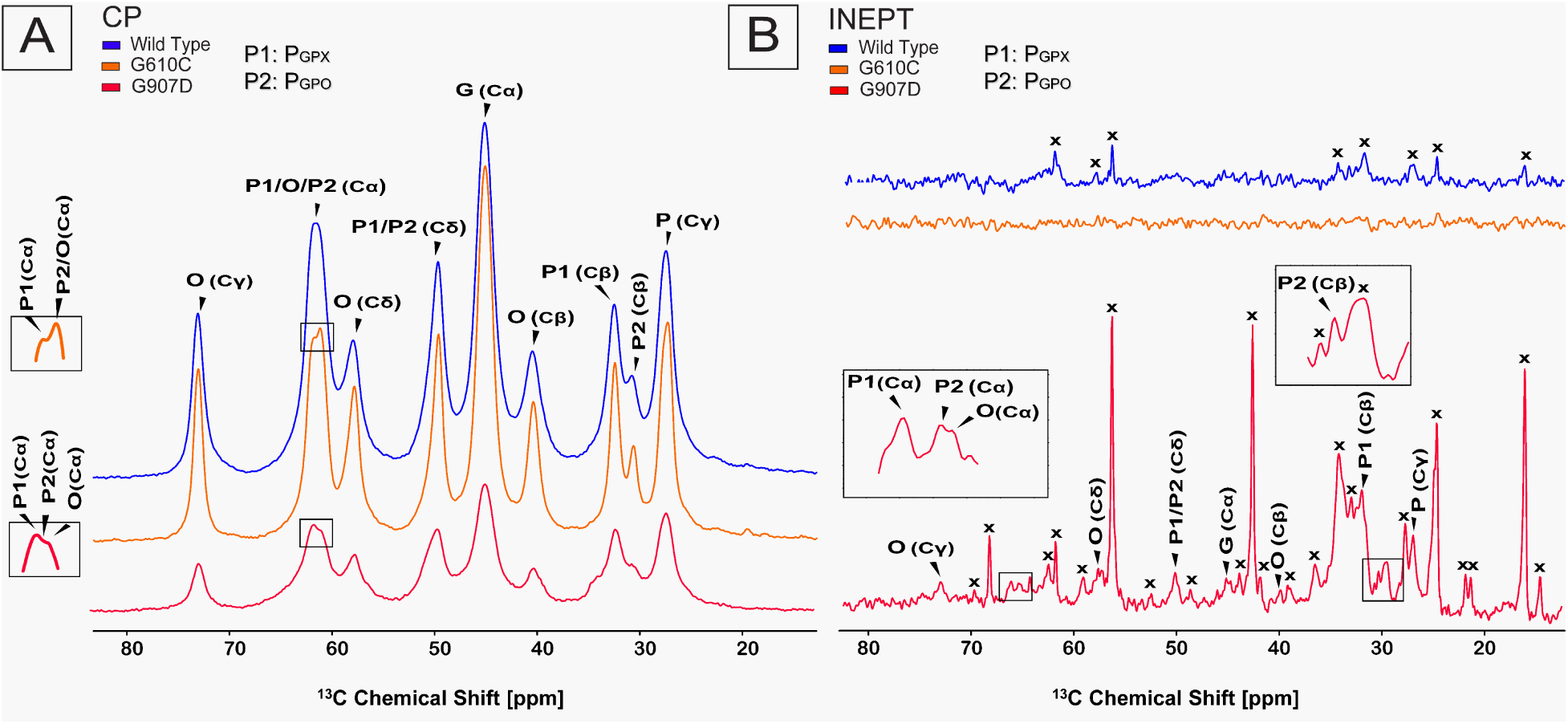
1D 13C-ssNMR of ECM variants. (**A**) 1H-13C CP spectra for WT (blue), G610C (orange), and G907D (red) ECM. The signal intensity of G907D is ∼2.6-fold lower than WT and 2.75-fold lower than G610C. Boxed regions show zoomed views of Pro and Hyp Cα peaks to highlight resolution between peaks. (**B**) 1H-13C INEPT spectra of WT (blue), G610C (orange), and G907D (red). Boxed regions show zoomed views of Pro and Hyp peaks to highlight resolution between peaks. Peaks unrelated to glycine, proline, and hydroxyproline are shown with a cross (x).

### Functional impact of OI mutaions on collagen-integrin interactions and cellular responses

We addressed the functional consequences of OI-associated mutations at both the protein and cellular levels by examining their effects on collagen-integrin interactions and ECM-mediated cellular responses. To assess this, at the protein level, enzyme linked immunosorbent assays (ELISA) were performed using the α2 I-domain of the α2β1 integrin, which contains the primary ligand binding site and has been shown to retain both the specificity and affinity of full-length α2β1 for its collagen ligands^17,20,43^. As such, it serves as a widely accepted model for studying α2β1–collagen interactions in structural and biochemical assays. Integrin–collagen binding depends on the coordination of a divalent metal cation at the metal ion-dependent adhesion site (MIDAS) within the I-domain. Therefore, all experiments were conducted in the presence of Mg²⁺ to support stable and physiologically relevant binding. The concentration-dependent binding of integrin to collagen fibrils revealed that G610C fibrils exhibit reduced binding affinity compared to both wild-type (WT) and G907D fibrils (Fig. 4). ELISA results further showed that integrin binds WT and G907D fibrils with similar affinity at lower concentrations, but G907D displays increased binding at the highest integrin concentration tested (20 µg/ml).

**Figure 4:**
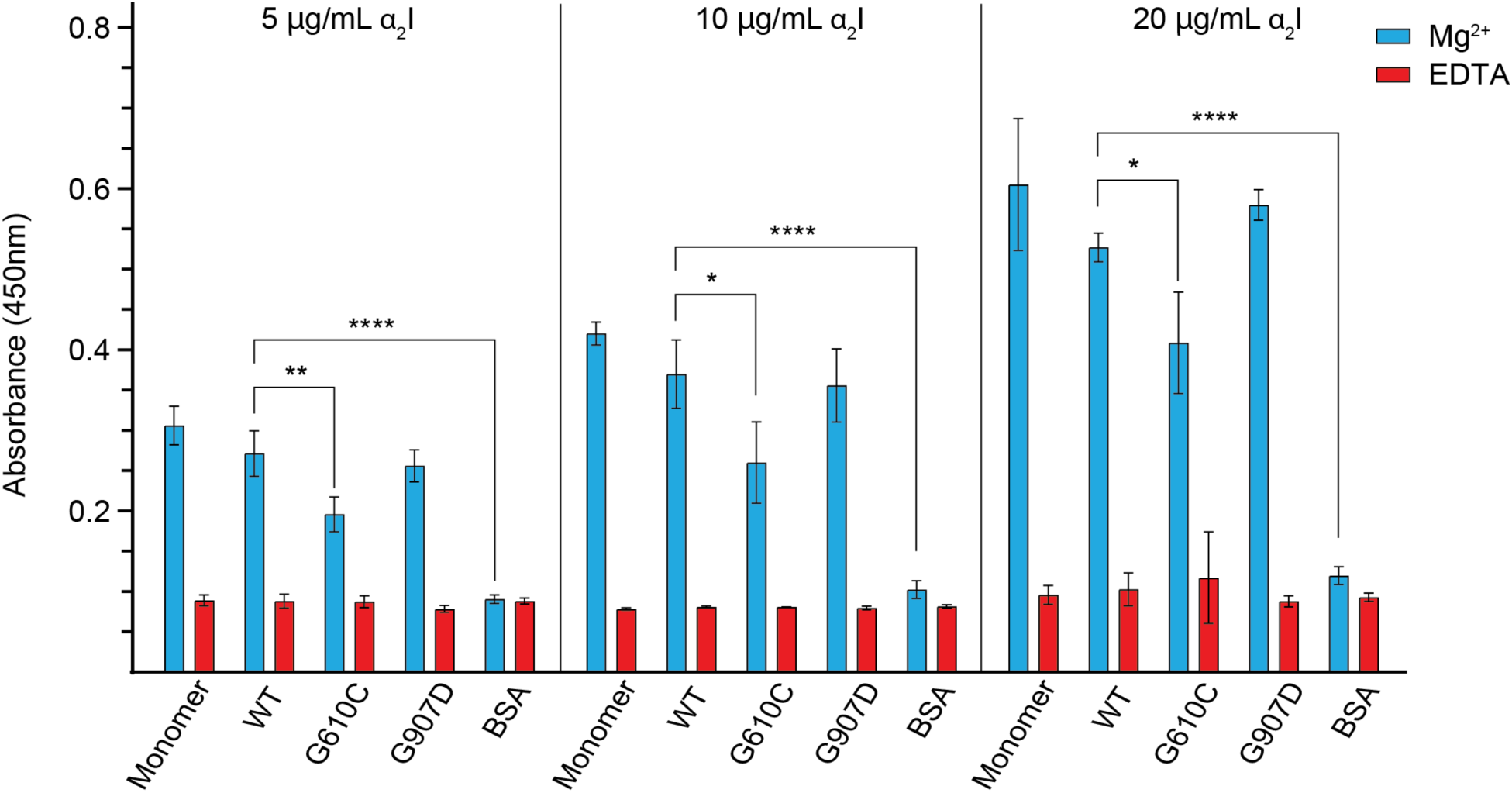
Dose-dependent binding of α_2_I-domain to primary human fibroblast collagen fibrils. ELISA binding assay shows collagen-integrin binding increases with concentrations of α_2_I-domain. Monomeric rat tail collagen I (Monomer), WT sonicated fibrils (WT), G610C sonicated fibrils (G610C), G907D sonicated fibrils (G907D), and BSA as negative control were used to determine the dose dependence of the collagen-integrin binding. All fibrils were in PBS before coating the plate. Monomeric collagen I was in 10mM acetic acid to retain triple helical properties while coating the plate. ELISA binding was performed in PBS with the addition of either 5mM MgCl_2_ (blue) or 5mM EDTA (red). Error bars indicate the standard deviation of the measurements, taken in quadruplicate. The asterisk (*) notations indicate a significant difference between the WT fibrils and the groups containing the notation. The “*” represent a *p*-value of <0.05, <0.01, <0.001, and <0.0001, respectively.

To directly visualize how integrins interact with WT and mutant collagen fibrils, we imaged collagen–α2 I-domain complexes using AFM. In phase images, integrin appears as dark, globular features on the fibril surface (Fig. 5A–D and Supplementary Fig S6). WT fibrils show prominent integrin binding (Fig. 5A), whereas G610C fibrils display significantly reduced integrin association (Fig. 5B). In contrast, G907D fibrils exhibit enhanced integrin binding, often accompanied by large, bulge-like structures attached to the fibrils (Fig. 5C). As expected, the negative control lacking Mg²⁺ shows minimal integrin association with WT fibrils (Fig. 5D). Additional representative AFM images are provided in Supplementary Fig. S6. Quantification of integrin features across fibril types (Fig. 5E and Supplementary Table S4) confirms that G610C reduces, while G907D increases, integrin binding relative to WT. The AFM results are consistent with ELISA-based binding assays, both showing reduced integrin binding in G610C and enhanced binding in G907D. Together with our TEM, ELISA and ssNMR data, these findings demonstrate that OI-linked mutations can alter both the nanoscale structure and integrin-binding capacity of collagen fibrils.

**Figure 5:**
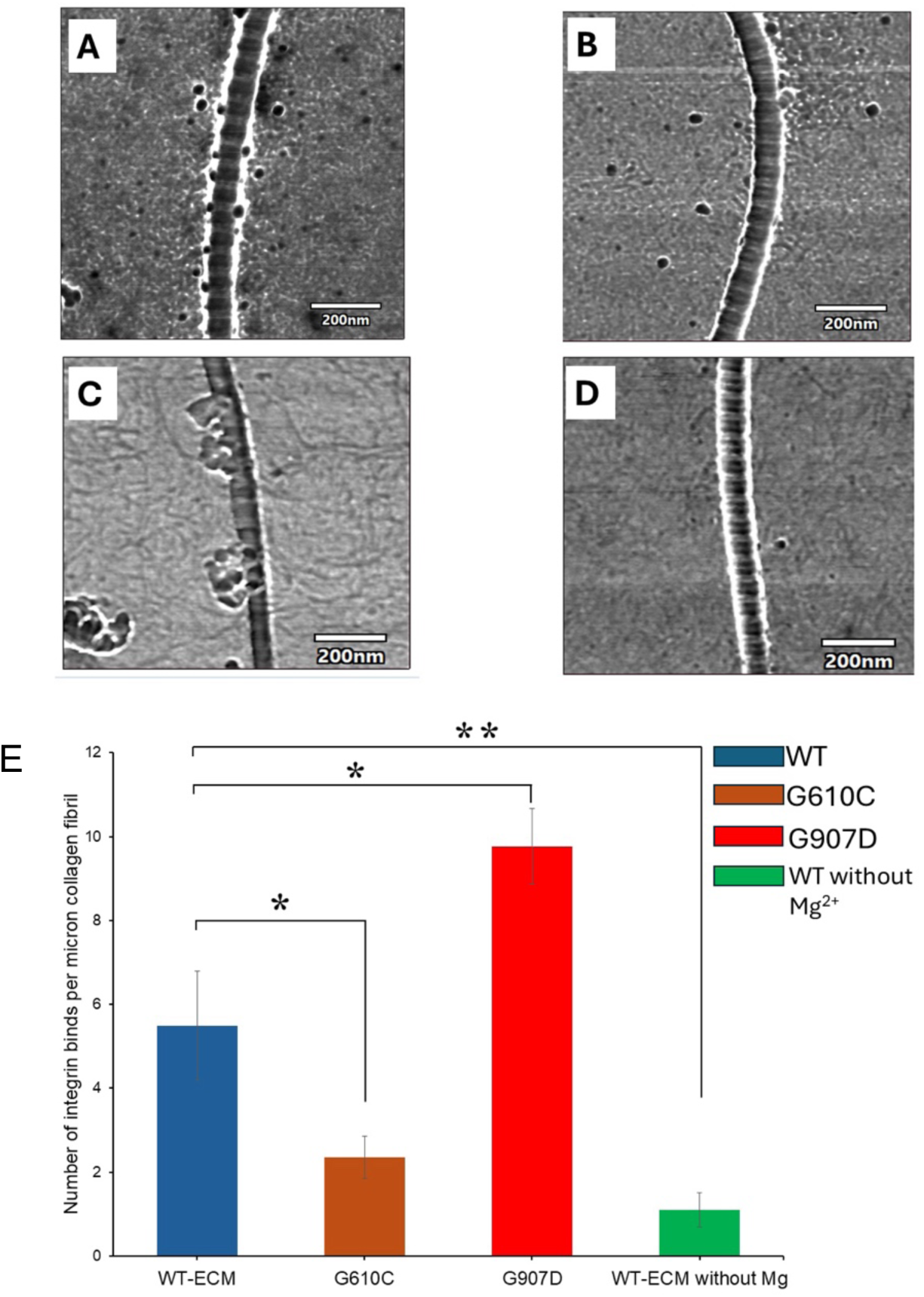
Direct visualization of α_2_I-domain binding to human fibroblast collagen fibrils. AFM phase images show the complex between α2I-domain and (A) WT ECM fibril, (B) G610C fibril, (C) G907D fibril, and (D) WT-ECM without Mg^2+^. (E) Bar plot compares the binding affinity of α_2_I integrin to WT-ECM and mutant fibrils. The symbols “*” and “**” represent p-value of <0.05 and <0.01 respectively.

To determine whether OI-associated collagen mutations alter broader ECM properties and cellular responses, we performed cell adhesion and proliferation assays using ECM derived from healthy donor fibroblasts (WT) and patient-derived fibroblasts carrying G610C (moderate OI) and G907D (lethal OI) mutations. MC3T3-E1 cells exhibited progressive proliferation over five days on all ECM substrates, with no significant differences observed between WT and mutant ECMs (Fig. 6). Similarly, cell adhesion was comparable across WT, G610C, and G907D ECMs (Fig 6). These results indicate that although mutation-induced nanoscale alterations in collagen fibrils can modulate collagen–integrin interactions, as detected by ELISA, they are insufficient to produce measurable changes in overall cell attachment or proliferation. While ELISA assesses direct protein-protein interactions, the adhesion assay primarily reflects overall cell retention and does not fully distinguish specific integrin-mediated adhesion from broader cellular responses. In addition, proliferation assays provide an indirect readout of integrated metabolic activity and growth, which may require more pronounced changes in ECM composition or organization to reveal detectable differences. Collectively, these findings suggest that OI-associated nanoscale fibrillar defects contribute to disease pathology, but that the severity of OI likely arises from cumulative effects on matrix organization, mineralization, and tissue remodeling during later stages of tissue development.

**Figure 6:**
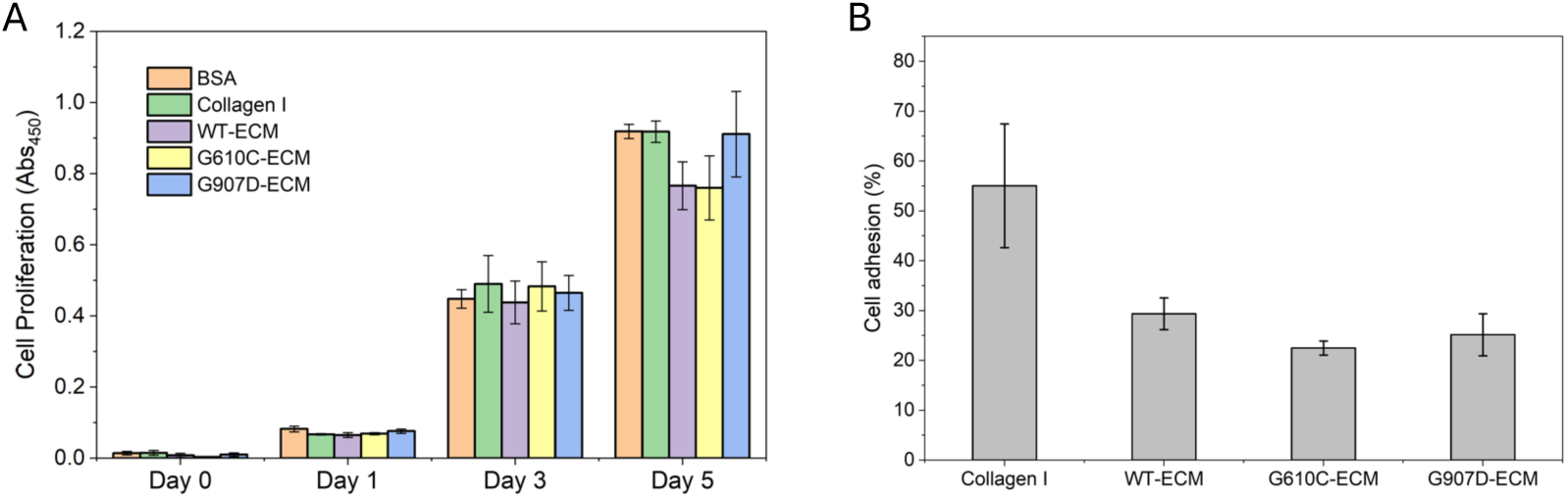
Cell proliferation and adhesion of MC3T3-E1 cells on ECM derived from wild-type and OI patient fibroblasts. (A) MC3T3-E1 cell proliferation was assessed using the Cell Counting Kit-8 (CCK-8) assay on surfaces coated with fibroblast-derived ECM from WT, G610C, and G907D samples. Cell proliferation was monitored over a 5-day period (Days 0, 1, 3, and 5) and expressed as absorbance at 450 nm. MC3T3-E1 cells exhibited progressive increases in proliferation across all ECM conditions, with no significant differences observed between WT, G610C, and G907D ECMs. Data are presented as mean ± SD; (B) MC3T3-E1 cell adhesion to the same coated surfaces was quantified using the CCK-8 assay. Comparable levels of cell adhesion were observed among WT, G610C, and G907D-derived ECMs, indicating that the OI-associated collagen mutations did not markedly affect overall cellular attachment. Data are presented as mean ± SEM. No significant differences in adhesion were observed among WT, G610C, and G907D ECMs.

### Differential lethality of collagen I α1 and α2 mutations in OI

We reinvestigated pathogenic collagen type I α1 and α2 genetic variants and associated OI Gly missense mutations to understand how G610C and G907D mutants relate to the greater landscape of OI mutations and to determine whether patterns of disease phenotype have changed or emerged since previous bioinformatic analyses of OI patient data^8,44^. In particular, the latest major bioinformatics study of OI collagen mutations was conducted by Marini et al. in 2022^8^ and focused on the impact of Gly missense and splice site mutations. The study reported 793 Gly substitutions for COL1A1 (including 241 lethal OI cases) and 709 Gly substitutions for COL1A2 (including 133 lethal OI cases). In the present study utilizing OI patient data from the Leiden Open Variation Database in 2025, we extracted a total of 986 Gly mutations (238 lethal OI cases) in the α1 chain and 924 Gly missense mutations (135 lethal OI cases) in the α2 chain (Fig. 7 and Supplementary Fig. S7), likely representing different data cleaning approaches between studies and an uptick in the number of reported nonlethal OI cases. The type of amino acid substitution clearly influences the phenotype in α1 and α2 chains (Fig. 7A and 7C). Regardless of alpha chain, mutations with small side chains are associated with lower lethality (α2-α1%: Ala 0.0-10.0%, Ser 7.1-14.6%, Cys 13.5-29.6%), whereas bulkier residues correspond to higher lethality (α2-α1%: Asp 23.9-51.2%, Glu 27.5-43.5%, Val 26.2-48.7%), in agreement with previous studies^10,27,45,46^. These observations support the view that larger amino acid substitutions introduce greater steric hindrance within the collagen triple helix, resulting in increased structural destabilization and a higher likelihood of severe or lethal OI phenotypes. It is also evident from this data that mutations in the α1 chain are more deleterious than those found in an α2 chain supporting previous data^9,44^. Odds ratio analysis of collagen I mutations^44^ shows that α1 mutations are twice as lethal as α2 mutations (Supplementary Table S5 and S6). Interestingly, mutation frequency varies across the alpha chains (Fig. 7B and 7D) with several hotspot locations showing noticeably more mutations, however the lethal G907 mutation does not fall in a high frequency region. Taken together, the findings from the present updated and expanded analysis provide further evidence reinforcing previously observed relationships between sequence and OI phenotype

**Figure 7:**
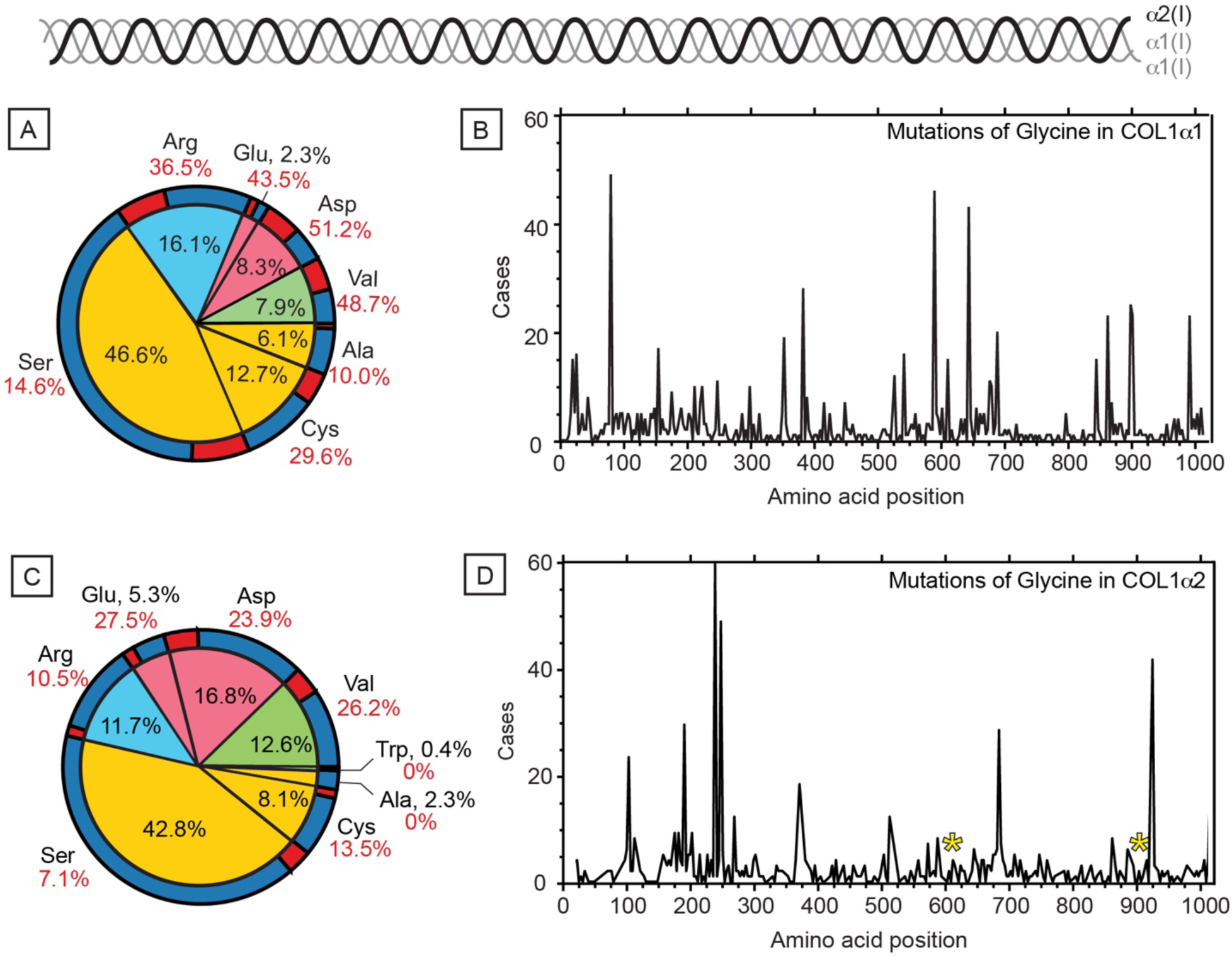
Comparison of OI Gly missense mutations in collagen I α1 and collagen I α2 chains. Summary analysis of Gly ◊ X missense mutations on the collagen I triple helix leading to OI, with data collected from the Leiden Open Variation Database. Blue, red, green, and yellow pie chart slices correspond to Gly mutations’ amino acid type (positively charged, blue; negatively charged, red; hydrophobic, green; small, yellow). Inner pie chart rings show overall percentages, and outer rings are subdivided based on lethality with respect to the specific amino acid substitution. **(A)** Percentages of Gly ◊ X amino acid mutations on collagen I α1. **(B)** Frequency chart of collagen I α1 chain mutations. **(C)** Percentages of Gly ◊ X amino acid mutations on collagen I α2. **(D)** Frequency chart of collagen I α2 chain mutations. Yellow asterisks represent the location of the studied G610C and G907D mutations.

## Discussion

Collagen I fibrils form the primary tensile framework of connective tissues, and alterations to their nanoscale structure can profoundly affect tissue mechanics, matrix–cell interactions, and disease phenotype. Osteogenesis imperfecta (OI) is a heritable connective tissue disorder most commonly caused by dominant mutations in the genes encoding α1 or α2 chains of collagen I. The majority of pathogenic mutants involve substitutions of a single glycine residue in the GXY repeating motif of the collagen triple helix. While several studies have investigated the role of mutations on the structure and dynamics of the collagen triple helix using model collagen mimetic peptides^13-15,37,47-49^, much remains unknown how these mutations propagate to, and affect, fibril structural integrity and function. While α1 chain mutations are generally associated with more severe phenotypes, pathogenic variants in the α2 chain can also produce the full clinical spectrum, including severe and perinatal lethal forms of OI (Fig. 7). These genotype– phenotype relationships highlight the challenges in predicting function and disease severity.

In the present study, we investigate the structural and functional characteristics of collagen fibrils derived from primary fibroblast ECM obtained from both healthy individuals and patients with OI. We focused on two different α2 chain glycine mutations with distinct clinical outcomes: G610C, associated with moderate severity, and G907D, linked to perinatal lethality. Collagen fibrils harvested from these fibroblast cultures were examined for nanoscale organization, supramolecular morphology, integrin-binding affinities and cell-based studies. By integrating nanoscale morphological analysis (AFM, TEM) with high-resolution conformational studies (solid-state NMR), and functional binding assays (ELISA and cell-based) with bioinformatics, we directly relate OI clinical phenotype to mutation-specific fibril structural defects and altered integrin-mediated interactions. Although dermal fibroblast-derived ECM does not recapitulate the tissue-specific environment of bone, including osteoblast-driven mineralization and matrix remodeling, fibroblasts remain a well-established model for studying type I collagen biosynthesis, fibrillogenesis, and the molecular consequences of OI-associated collagen mutations^50-52^. Therefore, our findings primarily provide insight into early events in collagen assembly and ECM organization that precede later stages of bone development and remodeling. While fibroblast derived ECM also contains other fibrillar collagens, including types III and V, type I collagen remains the predominant collagen species, and the mutation-specific differences observed here are interpreted primarily in the context of COL1A2 associated alterations in type I collagen.

Both G610C and G907D fibrils retain the hallmark D-banding pattern of WT collagen I, however their structural and functional properties differ measurably. G610C fibrils are morphologically similar to WT by AFM and TEM. This observation is consistent with previous studies of the Col1a2+/G610C mouse model, which demonstrated that bone morphology, mineral composition, and collagen cross-linking remain largely unchanged during early developmental stages but become progressively altered with age and mineralization^31,53^. Specifically, Masci et al. reported minimal structural differences between WT and G610C bone at 10 days of age, whereas significant changes emerged after two months, coinciding with the progression of mineralization. Our findings using ECM derived from patient fibroblasts may therefore reflect an early, pre-mineralization stage of collagen organization, during which fibril morphology remains relatively preserved despite the presence of the mutation. This interpretation suggests that the pathological consequences of the G610C mutation arise progressively through later events such as matrix remodeling and mineralization rather than through pronounced early disruptions in fibril architecture. By contrast, G907D fibrils display clear nanoscale defects, including overall reduced height, and localized fibril thinning features. Solid-state NMR reveals that these morphological abnormalities are accompanied by increased molecular mobility, including partial loss of polyproline II structure in the mobile regions. The differing effects of G610C and G907D further suggest that side-chain size alone does not fully determine the structural consequences of Gly substitutions. Although Cys and Asp possess side chains of comparable sizes, the G907D mutation occurs closer to the C-terminal region of the collagen triple helix, where helix nucleation is initiated and folding proceeds in a C-to-N direction^54^. Disruptions in this region are therefore expected to have a greater impact on triple-helical stability and fibril organization. We propose that G907D alters the collagen fibril in a spatially restricted manner, creating regions of abnormal flexibility and altered surface topology. In addition to altered fibril assembly, the reduced fibril height observed for G907D may also reflect partial intracellular retention and reduced secretion of mutant collagen molecules, a mechanism that has been reported for several OI-associated glycine substitutions within the triple-helical domain^5,55,56^. Although collagen secretion was not directly quantified in the present study, the biochemical nature of the Gly substitution is consistent with mutations known to impair collagen folding and trafficking through the secretory pathway. These findings are consistent with the phenotype of OI, where non-lethal cases display fibril morphology very similar to WT or healthy controls, whereas lethal cases exhibit comparatively more pronounced fibril damage. These features are further supported by our and previous bioinformatics analyses, which indicates that glycine substitutions with larger amino acids, for example, Gly to Asp, are associated with a higher frequency of lethal cases compared to substitutions with smaller residues such as Cys, regardless of whether the mutation occurs in the α1 or α2 chains.

When considering the morphological impact of single Gly->X mutations, it is important to consider that OI mutations are heterozygously expressed on only one allele. Because type I collagen contains two α1(I) chains and one α2(I) chain, a heterozygous *COL1A1* mutation produces mutant α1 chains half the time, resulting in 75% abnormal collagen triple helices, whereas a heterozygous *COL1A2* mutation affects only the single α2 chain, yielding 50% abnormal collagen triple helices^57^. In our present study, both G610C and G907D mutations occur in the α2 chain, and therefore we expect that only half of the α2 collagen chains will carry the mutation (Supplementary Fig. S7), while the other half will be wild-type. However, it remains unclear how these mutated triple helices are distributed across the fibrils—whether the distribution is even, random, or follows a specific pattern. This differential distribution of WT and mutated helices in the fibril could influence the assembly process and affect overall fibril morphology as well. One possible explanation for the retention of D-periodicity in G610C and G907D, despite the presence of mutations, is that WT and mutant triple helical monomers co-assemble within the same fibril, preserving the register of the D-banding even if local structural defects are present. However, the extent to which mutant triple helices are incorporated into individual fibrils remains unresolved and represents an important area for future investigation. Nevertheless, our results demonstrate that specific mutations can differentially affect the extent and nature of morphological disruption underscoring how even a single Gly mutation within the α2(I) chain can propagate along the triple helix to affect fibril morphology and dynamics.

In addition to determining nanoscale morphology in lethal and non-lethal OI, we next examined how mutations affect collagen I fibril interactions with integrin binding sites. Integrins recognize GXX′GEX′′ motifs within the triple-helical domains of collagen I via their αI-domains in a divalent cation–dependent manner, with effective binding requiring precise spatial presentation of these motifs on the fibril surface. Our data reveal that G610C fibrils exhibit significantly reduced integrin binding relative to both WT and G907D, despite retaining canonical D-banding and overall fibril morphology. Previous work has shown that glycine substitutions induce local triple-helix unfolding, thereby disturbing the conformational requirements for integrin binding^15^. We hypothesize that the proximity of G610C to the critical GFOGER-binding site within the D3 region may perturb the positioning of the essential Glu residue required for integrin recognition (Fig 1). Furthermore, several other integrin-binding GXX’-GEX” motifs (for instance GROGER, GQRGER, GASGER, GLOGER) neighbouring G610C (Supplementary Fig. S10),, may also lose their ability to bind integrin, further reducing overall binding. Previous studies using collagen-mimetic peptides have demonstrated that OI-associated glycine substitutions induce local kinks, altered triple-helical dynamics, disrupted integrin-mediated interactions^44,58,59^. Previous studies have shown that Gly→Cys substitutions in procollagen can generate conformational perturbations extending more than 200 nm from the mutation site^60^. Together, these findings suggest that the reduced integrin binding observed for G610C may arise from both local disruption of integrin-binding motifs and from propagates mutation-induced conformational changes within the collagen fibril that affect integrin binding.

In contrast, the G907D mutation, which introduces local nanoscale structural defects and dynamics, markedly enhances integrin binding. At higher receptor concentrations, G907D fibrils display frequent αI-domain clustering along the fibril surface. Previous studies by Baldwin et al.^61^ demonstrated that the G907D substitution in the α2(I) chain reduces the thermal stability of type I procollagen, indicating destabilization of the collagen triple helix. This reduced thermal stability is consistent with our observations of increased molecular mobility by ssNMR, localized structural changes and reduced fibril height detected by AFM. Together, these findings suggest that the G907D mutation compromises molecular packing and alters the supramolecular organization of collagen fibrils. Previous molecular dynamics simulations of WT fibrils have shown that normally buried integrin-binding motifs may be intermittently exposed via transient nanosecond-scale fluctuations^22^ thereby providing access to hidden binding sites. The combination of reduced triple-helical stability and enhanced fibril dynamics observed for G907D may therefore increase the accessibility of cryptic integrin-binding motifs. Our AFM and NMR experiments reveal enhanced structural deformation and mobility in the G907D mutant, suggesting that these changes may potentially expose otherwise hidden binding sites, promoting greater integrin accessibility and facilitating receptor clustering. Together, these results highlight how distinct glycine substitutions can differentially modulate integrin–collagen interactions: we propose that G610C impairs binding by destabilizing local triple helical conformation at integrin binding site, while G907D increases binding by exposing cryptic sites through nanoscale fibril deformation. Importantly, although sonication was required to disperse the collagen, identical conditions were applied to all samples. Moreover, the increased molecular mobility observed for G907D by ssNMR using non-sonicated samples supports an intrinsic structural and dynamic alteration rather than a consequence arising primarily from sonication-induced enrichment of monomeric collagen. These findings also underscore that preserved fibrillar banding alone is not a sufficient marker of functional consequences or disease severity. In summary, our integrated structural, dynamic, and functional analyses show that nanoscale collagen fibril behavior is more nuanced than morphology alone can reveal, and that mutations outside canonical binding motifs can still profoundly influence receptor engagement. Although no significant differences were observed in MC3T3-E1 cell adhesion and proliferation on WT, G610C, and G907D ECMs under homeostatic culture conditions, collagen-integrin interactions remain critical regulators of cellular responses during tissue remodeling, wound healing, and repair processes. Alterations in collagen receptor accessibility or binding affinity may potentially have more pronounced consequences under dynamic physiological or pathological conditions where integrin-mediated signaling is actively engaged.

Three principles emerge from our study: First, collagen fibrils with preserved average D-banding can still exhibit substantial functional impairment, underscoring the importance of integrating structural imaging with dynamic and biochemical analyses. Second, pathogeneic effects on functional binding can arise from mutations outside the canonical interaction motifs, as structural and dynamic alterations may propagate through the triple helix to remodel the accessibility of functional domains. Third, our integrated approach, linking nanoscale morphology, molecular dynamics^22^, and receptor-binding behavior, offers a broadly applicable framework for investigating pathogenic variants in other collagen systems, including those arising from aging and PTMs. Applying this multimodal approach across collagen-associated conditions may uncover structure–function relationships with translational relevance. Beyond OI, these findings may reveal general principles of collagen biology that extend across fibrillar types. Fibril-level disruptions can arise from diverse molecular defects in collagen, including aberrant post-translational modifications (PTMs), non-enzymatic glycation, or age-related oxidative damage, each of which can destabilize the triple helix, impair fibrillogenesis, and alter receptor engagement^62,63^. We also acknowledge that factors such as PTMs that alter surface charge could lead to changes in collagen microenvironments and may contribute to altered integrin accessibility. By linking mutation-specific structural perturbations to changes in biomechanical and cell-matrix interactions, this work provides a conceptual framework for understanding how both genetic and acquired perturbations to collagen contribute to connective tissue pathologies, and it lays the foundation for the rational design of targeted therapeutic strategies for OI and related disorders.

## Methods

### Cell culture for human fibroblasts

Human fibroblast primary cell lines GM17416 (*Col1a2*^+/+^, “WT”), GM17596 (*Col1a2*^+/G610C^, “G610C”), and GM10503 (*Col1a2*^+/G907D^, “G907D”) were purchased from the Coriell Institute (Camden, NJ). WT and G610C cell lines chosen were from adult biological female donors whose ages were within 5 years apart. The G907D primary cell donor was a perinatal biological female. The cell lines were purchased to create as little variation as possible, although the age of the G907D line could not be matched due to the lethality of the mutation. The mutations represent Gly^700^ and Gly^997^ in the Col1α2 FASTA sequence for the G610C and G907D mutations, respectively. Sequences of both collagen α-chains were checked and sequenced via AZENTA/GeneWIZ NGS Whole Exome sequencing (Fig. S8)^64^. WT and G610C cells were maintained with Eagle’s MEM with Earle’s Salts, non-essential amino acids (Millipore Sigma St. Louis, MO) supplemented with 2mM L-glutamine, 15% non-heat activated FBS (Gibco, Thermo Fisher Scientific, Inc., Waltham, MA), and 1% Penicillin/Streptomycin (Gibco, Thermo Fisher Scientific, Inc., Waltham, MA). G907D cells were maintained with Dulbecco’s Modified Eagle Medium (Gibco, Thermo Fisher Scientific, Inc., Waltham, MA) supplemented with 2mM L-glutamine, 10% non-heat activated FBS (Gibco, Thermo Fisher Scientific, Inc., Waltham, MA), and 1% Penicilllin/Streptomycin (Gibco, Thermo Fisher Scientific, Inc., Waltham, MA). All primary cells were maintained in a 37°C and 5% CO_2_ incubator. All cells and matrices for these experiments were grown in 75cm^2^ flasks and underwent media exchange every two to three days with their respective media.

### *In vitro* collagenous extracellular matrix expression from primary fibroblasts

All mammalian cell culture procedures for producing and extracting *in vitro* ECM were adapted from protocols used in the Duer lab, and from Franco-Barraza’s ECM production protocol.^42,65,66^ At approximately 70% confluency, the fibroblast cells were supplemented with L-ascorbic acid 2-phosphate sesquimagnesium salt hydrate (Millipore Sigma St. Louis, MO), to induce collagen production and matrix formation. WT and G610C cells were supplemented with 75 µg/mL of the stable L-ascorbic acid, and G907D cells were given 100µg/mL of the stable L-ascorbic acid. The higher ascorbic acid concentration used for G907D cultures was primarily due to the reduced ECM formation observed in this mutant line. Without increasing the ascorbic acid concentration, it was difficult to maintain robust matrix deposition. The media for the cell lines were exchanged every two to three days with their respective medias containing L-ascorbic acid for a minimum of four weeks, and maximum of five weeks after L-ascorbic acid addition, approximately when the dense matrix produced by the cells began to peel off the surface of the culture flask. For reproducibility, matrices were expressed for 30-31 days after the L-ascorbic acid addition. If the matrix began peeling off the surface of the flask, no more than a third of the matrix was allowed to prematurely peel from the flask prior to extraction.

### Extracellular matrix extraction and cleaning

All buffers used were filtered with a 0.22 um filter prior to addition into cell cultures and extractions. Cultures containing extracellular matrix sheets were washed twice with 1X PBS pH 7.4 and either frozen in 5 mL of 1X PBS at -80°C for storage or extracted immediately with 5 mL of warmed extraction buffer (20 mM NH_4_OH, 0.5% TritonX-100 in PBS). The cells were incubated in the extraction buffer for 10 minutes in a 37°C incubator. After incubation, the cells were transferred into a centrifuge tube and an equivalent volume of 1X PBS was used to dilute the solution overnight. The next day, the ECM-cell mixtures were spun down at 1500 rpm for 3 minutes and washed twice with 1X PBS before being treated with warm DNA extraction buffer (10 mM Tris-HCl, 2.5 mM MgCl_2_, 0.1 mM CaCl_2_, 10 µg/mL DNAse I in dH_2_O) for 30 minutes in a 37°C incubator. The ECM extract was then washed twice with 1X PBS before storing in a -20°C freezer. All ECM sheet pellets were gently resuspended by inversion of the centrifuge tubes during washes and extractions to ensure the entire ECM sheet was washed by the buffer or exposed for treatments. If the flask was frozen prior to extraction, after thawing, the PBS was aspirated, and the ECM sheet was treated with warmed extraction buffer as described previously.

The ECM extracts were further processed using α-chymotrypsin (Millipore Sigma St. Louis, MO) to clean the non-collagenous ECM proteins from the surface of the fibrils. α-chymotrypsin was added using mass ratios, where “1X” α-chymotrypsin added is the recommended ratio of 1:60 α-chymotrypsin:ECM protein by mass. For all cell lines, we used a rough estimate of 24mg of ECM per 175cm^2^ surface area to estimate the mass of the ECM protein required for the weight ratio, based on prior *in vitro* lyophilized sample masses. The ECM was incubated with α-chymotrypsin buffer (100 mM Tris-HCl, 10 mM CaCl_2_) containing the predetermined amount of α-chymotrypsin at 37°C for 48 hours. For the TEM visualization, the samples were cleaned using a 3X concentration of the α-chymotrypsin for the 48 hours with gentle inversions. Inversions were done a total of 5 times, spread out over the span of 48 hours with a minimum of 4 hours between each inversion period, and at least 5 inversions of the tube during each step. After incubation, the sample was washed 3 times with 1X PBS to remove remaining α-chymotrypsin. For AFM, TEM, and ELISA experiments, ECM samples were treated with α-chymotrypsin to better resolve collagen fibrillar components, whereas NMR and cellular assays were performed using untreated ECM samples.

### Collagen fibril sample preparation

ECM was put into a 1.5 mL microcentrifuge tube and resuspended in 500 µL of PBS prior to sonication. The sample was sonicated using a tip sonicator (Fisher Sci Sonic Dismembrator Model 500) for 4s at 30% amplitude twice (8s total) with a minimum of 30s on ice between sonication steps and following the sonication. Any large fragments of ECM that remain after sonication were removed from the centrifuge tube as best as possible and remaining fibril solutions were kept on ice or stored at 4°C until used in experiments. Larger fragments that were unable to be removed were avoided to keep samples as uniform as possible. Additionally, the large pieces/sections of ECM were avoided for TEM/AFM as they could not be visualized by either technique due to the thickness of the sheets. For similar reasons, unsonicated samples were not imaged.

To determine the approximate concentration of collagen, sonicated fibrils were run on an SDS-PAGE gel and compared to lanes with bottled rat tail collagen. (Supplementary Fig. S9) Due to the inherently random nature of fibril sizes after sonication, concentrations of the sonicated collagen fibrils can only be estimated for assays. However, the same preparation was used for all fibrils to reduce the potential for errors. For SDS-PAGE, the fibril samples were loaded with a final concentration of 1X Laemmli buffer (4X solution, BioRad Hercules, CA) with 355 mM β-mercaptoethanol (BioRad Hercules, CA). The samples were thermally denatured at 85°C for 15 minutes in the loading buffer just prior to running the gel. The samples were analyzed by SDS-PAGE 4-15% gradient polyacrylamide gels (BioRad Hercules, CA) and run in a Mini Protean Tetra Cell chamber. The gels were run at 50V for 5 minutes, followed by 100V for 60 minutes. Gels were fixed (40% EtOH, 10% Acetic Acid, 50% water) for 15 minutes, and stained with QC Coomassie Colloidal Blue (BioRad Hercules, CA)^67^ for 18 to 20 hours. Destaining was done with ultrapure water for 2 hours, changing the water a minimum of 3 times. Fixation, staining, and destaining steps were all done at room temperature on a gel rocker at 50rpm, and images were taken immediately after destaining.

### AFM experimentation and Analysis

50 µL of ECM suspension was placed onto a freshly cleaved mica surface and allowed to bind for 5 minutes before being washed with 2 mL of ultrapure (Millipore) water. The sample was then allowed to dry for 1 hour in a laminar flow hood. Afterward, the sample was placed into the AFM for imaging. All imaging was performed on a Cypher ES AFM (Asylum Research) with AC240 tips (nominal resonance frequency of 70 kH, nominal spring constant of 2 N/m) and at room temperature. For analysis, height data from the images were imported into MATLAB and homemade scripts were used to determine the height and periodicity of the fibril samples.

### TEM experimentation

200/300 mesh Cu carbon coated grids (Electron Microscopy Sciences Hatfield, PA) were used for TEM experiments. 5 µL of the sample were placed on the grid and incubated at room temperature for 2 minutes before removal of the remaining liquid by blotting the side of the grid onto filter paper. The grids were negatively stained for 1 minute using 3 µL of 3% uranyl acetate (Electron Microscopy Sciences Hatfield, PA). Excess liquid was blotted off and allowed to dry. The TEM grids were dried at least overnight before imaging, preferably for at least 48h.TEM images were performed on a JEOL JEM 2010F microscope at 200.0kV. Bright-field transmission electron microscopy (BF-TEM) images were analyzed with ImageJ software. From densitometry graphs regional light intensity was used to measure fibril width of genetic variants consisting of wild type (WT), G610C (mild OI), and G907D (lethal OI). An ANOVA one way test was performed followed by a Tukey test to determine significance between types.

### Statistical Analysis of AFM data

GraphPad PRISM software was used in the statistical analysis of the AFM data. One-way ANOVA with multiple comparison tests were run for the WT, G610C, and G907D samples. The statistical tests ran for the samples assumed gaussian distribution for all samples and non-equivalent standard deviations between samples.

### NMR

By enriching cell cultures with ascorbic acid and U-^13^C,^15^N-Gly, and U-^13^C,^15^N-Pro amino acids, we produced isotopically labeled collagen fibrils in *in vitro* ECM from fibroblast cell line and investigated the structural and dynamic characteristics of these fibrils using ssNMR spectroscopy. We isotopically labeled the ECM samples with Gly and Pro because these residues are the most abundant amino acids within the collagen I sequence and are primarily found in the triple helical region of the monomer and fibril structures. Due to the PTM of Pro in the third position of G-X-X’ triplets to Hyp, ^13^C,^15^N-Hyp was also present.

All ssNMR experiments were collected on a Bruker Avance III 600MHz (14.1T) solids magnet with a TXI probe at 25 °C and 11 kHz MAS. Isotopically labeled fibroblast ECM samples were lyophilized. Approximately 3.4 mg of material was packed into 1.6mm rotors, and rehydrated up to 60%. To measure rigid residues in collagen fibrils, 1D ^1^H-^13^C CP and 2D ^13^C-^13^C Dipolar Assisted Rotational Resonance (DARR) experiments were performed, which rely on 1H-13C cross-polarization to transfer magnetization onto insensitive 13C nuclei. Whereas mobile residues in the fibrils were detected with 1D ^13^C refocused Insensitive Nuclei Enhanced by Polarized Transfer (rINEPT) experiments, scalar J-coupling was used to transfer magnetization onto the ^13^C nuclei. NMR data were processed using Topspin 4.10 (Bruker), NMR Pipe/NMRDraw (Delaglio et al. 1995), and analyzed using Sparky (Lee et al. 2015) and CcpNmr Analysis.

### Integrin α2I domain expression and purification

Integrin α_2_I domain, which consists of residues 142-336 of WT α_2_ integrin, was recombinantly expressed using Escherichia Coli BL21(DE3) cells and incorporated an N-terminal His_6_ tag. Induction with 1 mM of IPTG for 16 hr at 25° C was employed to initiate protein production. Cells were then lysed using a 20% sucrose TES buffer. Next, the protein was purified using a Ni^2+^ -charged HisTrap HP column (GE Healthcare Life) and soon after buffer exchanged to PBS, pH 7.4, using a PD-10 desalting column (GE Healthcare Life).

### ELISA binding assays

ELISA assays were done similarly to our previous work.^24^ Immulon 2HB plates were coated in quadruplicate with the bottled collagen, WT sonicated fibrils, G610C sonicated fibrils, G907D sonicated fibrils, and PBS as a negative control overnight at 4°C. Bottled rat tail collagen (Corning, Edison, NJ) was diluted to 10 µg/mL in 20 mM acetic acid to ensure collagen adsorbed onto the plate was monomeric. All fibrils used in the assay were prepared as described previously. The concentration of the fibrils was diluted in PBS to an approximate concentration of 10 µg/mL (approximate concentration after denaturation). The coating solutions were discarded the following day and the wells being used, including the negative control were blocked with 200 µL of 50 mg/mL of BSA in PBS for 1h, rocking at 4°C. The wells were then given 100 µL of 5/10/20 µg/mL of α2I integrin diluted in PBS containing either 5 mM MgCl_2_ or 5mM EDTA. The integrin was incubated for 1h, followed by treatment with antibodies. 100 µL of mouse anti-α2I integrin primary antibody was added at a 1:2000 dilution for 45 minutes, followed by 100 µL of goat HRP-conjugated anti-mouse antibody at a 1:5000 dilution for 30 minutes. A TMB Substrate Kit (Pierce, Waltham, MA) was used according to the manufacturer protocol to measure the binding activity and absorbance was measured at 450 nm using an Infinite F50 microplate reader (TECAN, Männedorf, CH). Following each binding addition step, the solution(s) in the plates were discarded and the wells were washed thrice with the wash buffer (1 mg/mL BSA, 5 mM MgCl_2_/5 mM EDTA in PBS). Statistical significance between samples was measured via one-way ANOVA with multiple comparisons tests.

### Cell proliferation and adhesion assays

Cell proliferation and adhesion assays were performed as previously described^68^. 96-well tissue culture plates (Corning, USA) were coated overnight at 4 °C with the indicated substrates at a final concentration of 10 μg cm^-2^. Type I collagen was prepared in 0.02 N acetic acid, whereas ECM samples derived from WT, G610C, and G907D fibroblast cultures were diluted in PBS prior to coating. For the proliferation assay, wells were coated with BSA, type I collagen (Corning, USA) and ECMs derived from WT, G610C, and G907D fibroblast cultures. The sample solutions were removed and blocked by 5% BSA. MC3T3-E1 cells were seeded at a density of 2 x 10^3^ cells per well and cultured in α-MEM supplemented with 10% heat-inactivated fetal bovine serum and 1% penicillin-streptomycin at 37 °C in a humidified atmosphere containing 5% CO_2_. Cell proliferation was assessed on days 0, 1, 3, and 5 using the Cell Counting Kit-8 (CCK-8 from Sigma-Aldrich) assay according to the manufacturer’s instructions. Briefly, 10 μL of CCK-8 reagent was added to each well, and absorbance at 450 nm was measured using a microplate reader (FluoStar Omega, BMG Labtech). Culture medium was replaced every two days.

For the adhesion assay, plates were coated with type I collagen or ECM preparations derived from WT, G610C, and G907D fibroblast cultures. MC3T3-E1 cells were seeded at a density of 1 x 10^4^ cells per well and allowed to adhere for 6 h. Non-adherent cells were removed by gently washing the wells three times with PBS, and the number of adherent cells was quantified using the CCK-8 assay. Cell adhesion was expressed as the percentage of adherent cells relative to the initial number of seeded cells.

### AFM Analysis of Integrin α2I and Collagen Fibril Interaction

The interaction between Integrin α2I and collagen fibrils was studied using atomic force microscopy (AFM). For imaging integrin monomers as a control, 50 µL of Integrin α2I was placed on freshly cleaved mica surfaces. To observe the natural structure and mutation effects on ECM fibril D-banding, wild-type (WT), G610C, and G907D fibrils were separately imaged. All fibrils used in the assay were prepared as described previously. The fibrils were diluted in PBS to an approximate concentration of 125 µg/mL (calculated after denaturation).

To investigate integrin binding and alignment on fibril networks, Integrin α2I, collagen fibrils, and MgCl₂ were mixed in a 1:4:150 ratio and incubated for 1 hour at room temperature before AFM imaging. Since Mg²⁺ plays a key role in α2I-collagen binding, a control experiment was performed using samples containing integrin and WT fibrils (1:4 ratio) without MgCl_2_, incubated under the same conditions for 1 hour before imaging.

The number of binding sites where integrin interacted with collagen fibrils was analyzed using amplitude and phase AFM images.

### OI Mutation Database Analyses

OI mutation data was collected from the Leiden Open Variation Database version 3.0 (LOVD3).^12^ Mutations in the collagen protein sequences are described using the nomenclature guidelines created by the Human Genome Variation Society.^69,70^ For the database analyses, OI cases were categorized by disease lethality including lethal (type II) and nonlethal cases (types I, III, IV). Cases were determined by patient survival during the immediate postnatal or the prenatal period. Given the consistent appearance and structural importance of Gly in collagen I α1 and α2 chains, the bioinformatics analysis only contained Gly missense mutations to corroborate previous studies and further inform the relationship between collagen genotype and phenotype.

Python programming was used to extract data from LOVD3 text files. After removing redundant information from the dataset, a total of 986 cases of Gly missense mutations in the α1(I) chain and 924 cases in the α2(I) chain were included in the analysis. Since the most recent analysis by our group was conducted in 2016, there has been a significant increase in the number of cases and available data.^44^ The most recent search of the data was performed on June 19, 2025.

## Author Contributions

J.B designed the research; C.T., S.B harvested ECM; J.R, S.B., A. Nikhad performed and analyzed AFM experiments; P.R.F analyzed TEM data, A. Nayer, C.L.H designed and performed NMR experiments; C.T and A. Nikhad performed ELISA experiment; E.K.L performed bioinformatic analysis; S.B performed cell-based assays, S.B, C.L.H, J.R and J.B analyzed data; S.B, C.T, C.L.H, and J.B wrote the manuscript.

## Acknowledgements

This work was supported by an NIH grant GM136431 to Jean Baum and NIH T32 Postdoctoral Training Program in Translational Research in Regenerative Medicine under award number T32EB005583 to Jonathan Roth.

## References

1 Iozzo, R. V. & Gubbiotti, M. A. Extracellular matrix: The driving force of mammalian diseases. Matrix Biol 71-72, 1-9 (2018).

2 Hynes, R. O. The extracellular matrix: not just pretty fibrils. Science 326, 1216–1219 (2009).

3 Frantz, C., Stewart, K. M. & Weaver, V. M. The extracellular matrix at a glance. J Cell Sci 123, 4195–4200 (2010).

4 Sweeney, S. M. et al. Candidate cell and matrix interaction domains on the collagen fibril, the predominant protein of vertebrates. J Biol Chem 283, 21187–21197 (2008).

5 Jovanovic, M., Guterman-Ram, G. & Marini, J. C. Osteogenesis Imperfecta: Mechanisms and Signaling Pathways Connecting Classical and Rare OI Types. Endocr. Rev. 43, 61–90 (2022).

6 Claeys, L. et al. Collagen transport and related pathways in Osteogenesis Imperfecta. Hum Genet 140, 1121–1141 (2021).

7 Kuivaniemi, H., Tromp, G. & Prockop, D. J. Mutations in fibrillar collagens (types I, II, III, and XI), fibril-associated collagen (type IX), and network-forming collagen (type X) cause a spectrum of diseases of bone, cartilage, and blood vessels. Hum. Mutat. 9, 300–315 (1997).

8 Garibaldi, N. et al. Dissecting the phenotypic variability of osteogenesis imperfecta. Dis Model Mech 15 (2022).

9 Marini, J. C. et al. Consortium for osteogenesis imperfecta mutations in the helical domain of type I collagen: regions rich in lethal mutations align with collagen binding sites for integrins and proteoglycans. Hum. Mutat. 28, 209–221 (2007).

10 Qiu, Y. et al. Collagen Gly missense mutations: Effect of residue identity on collagen structure and integrin binding. J Struct Biol 203, 255–262 (2018).

11 Bregou Bourgeois, A., et al. Osteogenesis imperfecta: from diagnosis and multidisciplinary treatment to future perspectives. Swiss Med Wkly 146, w14322 (2016).

12 Fokkema, I. et al. The LOVD3 platform: efficient genome-wide sharing of genetic variants. Eur J Hum Genet 29, 1796–1803 (2021).

13 Xiao, J., Cheng, H., Silva, T., Baum, J. & Brodsky, B. Osteogenesis imperfecta missense mutations in collagen: structural consequences of a glycine to alanine replacement at a highly charged site. Biochemistry 50, 10771–10780 (2011).

14 Brodsky, B. & Baum, J. Structural biology: Modelling collagen diseases. Nature 453, 998–999 (2008).

15 Hoop, C. L. et al. Molecular underpinnings of integrin binding to collagen-mimetic peptides containing vascular Ehlers-Danlos syndrome-associated substitutions. J Biol Chem 294, 14442–14453 (2019).

16 Malcor, J. D. et al. Deciphering the folding code of collagens. Nat Commun 16, 2702 (2025).

17 Tuckwell, D., Calderwood, D. A., Green, L. J. & Humphries, M. J. Integrin alpha 2 I-domain is a binding site for collagens. J Cell Sci 108 (Pt 4), 1629–1637 (1995).

18 Adorno-Cruz, V. & Liu, H. Regulation and functions of integrin alpha2 in cell adhesion and disease. Genes Dis 6, 16–24 (2019).

19 Hamaia, S. & Farndale, R. W. Integrin recognition motifs in the human collagens. Adv Exp Med Biol 819, 127–142 (2014).

20 Knight, C. G. et al. The collagen-binding A-domains of integrins alpha(1)beta(1) and alpha(2)beta(1) recognize the same specific amino acid sequence, GFOGER, in native (triple-helical) collagens. J Biol Chem 275, 35–40 (2000).

21 Hoop, C. L., Zhu, J., Nunes, A. M., Case, D. A. & Baum, J. Revealing Accessibility of Cryptic Protein Binding Sites within the Functional Collagen Fibril. Biomolecules 7 (2017).

22 Zhu, J., Hoop, C. L., Case, D. A. & Baum, J. Cryptic binding sites become accessible through surface reconstruction of the type I collagen fibril. Sci Rep 8, 16646 (2018).

23 Roth, J., Hoop, C., Williams, J. K., Nanda, V. & Baum, J. Real-time single-molecule observation of incipient collagen fibrillogenesis and remodeling. Proc Natl Acad Sci U S A 121, e2401133121 (2024).

24 Roth, J., Hoop, C. L., Williams, J. K., Hayes, R. & Baum, J. Probing the effect of glycosaminoglycan depletion on integrin interactions with collagen I fibrils in the native extracellular matrix environment. Protein Sci 32, e4508 (2023).

25 Sarathchandra, P., Pope, F. M. & Ali, S. Y. An ultrastructural and immunogold localization study of proteoglycans associated with the osteocytes of fetal bone in osteogenesis imperfecta. Calcif Tissue Int 58, 435–442 (1996).

26 Baum, J. & Brodsky, B. Folding of peptide models of collagen and misfolding in disease. Curr Opin Struct Biol 9, 122–128 (1999).

27 Xiao, J., Madhan, B., Li, Y., Brodsky, B. & Baum, J. Osteogenesis imperfecta model peptides: incorporation of residues replacing Gly within a triple helix achieved by renucleation and local flexibility. Biophys J 101, 449–458 (2011).

28 Xu, K., Nowak, I., Kirchner, M. & Xu, Y. Recombinant collagen studies link the severe conformational changes induced by osteogenesis imperfecta mutations to the disruption of a set of interchain salt bridges. J Biol Chem 283, 34337–34344 (2008).

29 Fu, I., Case, D. A. & Baum, J. Dynamic Water-Mediated Hydrogen Bonding in a Collagen Model Peptide. Biochemistry 54, 6029–6037 (2015).

30 Li, Y., Brodsky, B. & Baum, J. NMR conformational and dynamic consequences of a gly to ser substitution in an osteogenesis imperfecta collagen model peptide. J Biol Chem 284, 20660–20667 (2009).

31 Masci, M. et al. Bone mineral properties in growing Col1a2(+/G610C) mice, an animal model of osteogenesis imperfecta. Bone 87, 120–129 (2016).

32 Kwon, J. & Cho, H. Collagen piezoelectricity in osteogenesis imperfecta and its role in intrafibrillar mineralization. Commun Biol 5, 1229 (2022).

33 Andriotis, O. G. et al. Structure-mechanics relationships of collagen fibrils in the osteogenesis imperfecta mouse model. J R Soc Interface 12, 20150701 (2015).

34 Sarathchandra, P., Pope, F. M., Kayser, M. V. & Ali, S. Y. A light and electron microscopic study of osteogenesis imperfecta bone samples, with reference to collagen chemistry and clinical phenotype. J Pathol 192, 385–395 (2000).

35 Traub, W., Arad, T., Vetter, U. & Weiner, S. Ultrastructural studies of bones from patients with osteogenesis imperfecta. Matrix Biol 14, 337–345 (1994).

36 Sarathchandra, P. & Pope, F. M. Unexpected ultrastructral changes in bone osteiod collagens in osteogenesis imperfecta. Micron 36, 696–702 (2005).

37 Hyde, T. J., Bryan, M. A., Brodsky, B. & Baum, J. Sequence dependence of renucleation after a Gly mutation in model collagen peptides. J Biol Chem 281, 36937–36943 (2006).

38 Bansode, S. et al. Glycation changes molecular organization and charge distribution in type I collagen fibrils. Sci Rep 10, 3397 (2020).

39 Chow, W. Y. et al. NMR spectroscopy of native and in vitro tissues implicates polyADP ribose in biomineralization. Science 344, 742–746 (2014).

40 De Sa Peixoto, P., Laurent, G., Azaïs, T. & Mosser, G. Solid-state NMR study reveals collagen I structural modifications of amino acid side chains upon fibrillogenesis. J Biol Chem 288, 7528–7535 (2013).

41 Saitô, H. et al. A high-resolution 13C-NMR study of collagenlike polypeptides and collagen fibrils in solid state studied by the cross-polarization-magic angle-spinning method. Manifestation of conformation-dependent 13C chemical shifts and application to conformational characterization. Biopolymers 23, 2279-2297 (1984).

42 Chow, W. Y. et al. Proline provides site-specific flexibility for in vivo collagen. Sci Rep 8, 13809 (2018).

43 Kamata, T. & Takada, Y. Direct binding of collagen to the I domain of integrin alpha 2 beta 1 (VLA-2, CD49b/CD29) in a divalent cation-independent manner. J Biol Chem 269, 26006-26010 (1994).

44 Xiao, J., Yang, Z., Sun, X., Addabbo, R. & Baum, J. Local amino acid sequence patterns dominate the heterogeneous phenotype for the collagen connective tissue disease Osteogenesis Imperfecta resulting from Gly mutations. J Struct Biol 192, 127–137 (2015).

45 Bhate, M., Wang, X., Baum, J. & Brodsky, B. Folding and conformational consequences of glycine to alanine replacements at different positions in a collagen model peptide. Biochemistry 41, 6539–6547 (2002).

46 Bodian, D. L., Madhan, B., Brodsky, B. & Klein, T. E. Predicting the clinical lethality of osteogenesis imperfecta from collagen glycine mutations. Biochemistry 47, 5424–5432 (2008).

47 Xu, Y. & Kirchner, M. Collagen Mimetic Peptides. Bioengineering (Basel) 8 (2021).

48 Fallas, J. A., Gauba, V. & Hartgerink, J. D. Solution structure of an ABC collagen heterotrimer reveals a single-register helix stabilized by electrostatic interactions. J Biol Chem 284, 26851–26859 (2009).

49 Yu, L. T. et al. Exploration of the hierarchical assembly space of collagen-like peptides beyond the triple helix. Nat Commun 15, 10385 (2024).

50 Marini, J. C. et al. Osteogenesis imperfecta. Nat Rev Dis Primers 3, 17052 (2017).

51 Pocock, A. E., Francis, M. J. & Smith, R. Type 1 collagen synthesis by skin fibroblasts from 17 patients with osteogenesis imperfecta type III. Clin Chim Acta 243, 53–72 (1995).

52 Brenner, R. E. et al. Altered collagen metabolism in osteogenesis imperfecta fibroblasts: a study on 33 patients with diverse forms. Eur J Clin Invest 20, 8–14 (1990).

53 Daley, E. et al. Variable bone fragility associated with an Amish COL1A2 variant and a knock-in mouse model. J. Bone Miner. Res. 25, 247–261 (2010).

54 Buevich, A. V., Silva, T., Brodsky, B. & Baum, J. Transformation of the mechanism of triple-helix peptide folding in the absence of a C-terminal nucleation domain and its implications for mutations in collagen disorders. J Biol Chem 279, 46890–46895 (2004).

55 Forlino, A., Cabral, W. A., Barnes, A. M. & Marini, J. C. New perspectives on osteogenesis imperfecta. Nat Rev Endocrinol 7, 540–557 (2011).

56 Lindert, U. et al. Insight into the Pathology of a COL1A1 Signal Peptide Heterozygous Mutation Leading to Severe Osteogenesis Imperfecta. Calcif. Tissue Int. 102, 373–379 (2018).

57 Gajko-Galicka, A. Mutations in type I collagen genes resulting in osteogenesis imperfecta in humans. Acta Biochim Pol 49, 433–441 (2002).

58 Xiao, J., Cheng, H., Silva, T., Baum, J. & Brodsky, B. Osteogenesis Imperfecta Missense Mutations in Collagen: Structural Consequences of a Glycine to Alanine Replacement at a Highly Charged Site. Biochemistry 50, 10771–10780 (2011).

59 Brodsky, B. & Baum, J. Modelling collagen diseases. Nature 453, 998–999 (2008).

60 Vogel, B. E. et al. A substitution of cysteine for glycine 748 of the alpha 1 chain produces a kink at this site in the procollagen I molecule and an altered N-proteinase cleavage site over 225 nm away. J Biol Chem 263, 19249–19255 (1988).

61 Baldwin, C. T., Constantinou, C. D., Dumars, K. W. & Prockop, D. J. A single base mutation that converts glycine 907 of the alpha 2(I) chain of type I procollagen to aspartate in a lethal variant of osteogenesis imperfecta. The single amino acid substitution near the carboxyl terminus destabilizes the whole triple helix. J Biol Chem 264, 3002–3006 (1989).

62 Sloseris, D. & Forde, N. R. AGEing of collagen: The effects of glycation on collagen’s stability, mechanics and assembly. Matrix Biol 135, 153–160 (2025).

63 Kirkness, M. W., Lehmann, K. & Forde, N. R. Mechanics and structural stability of the collagen triple helix. Curr Opin Chem Biol 53, 98–105 (2019).

64 Robinson, J. T. et al. Integrative genomics viewer. Nat Biotechnol 29, 24–26 (2011).

65 Franco-Barraza, J., Beacham, D. A., Amatangelo, M. D. & Cukierman, E. Preparation of Extracellular Matrices Produced by Cultured and Primary Fibroblasts. Curr Protoc Cell Biol 71, 10 19 11–10 19 34 (2016).

66 Muller, K. H. et al. Poly(ADP-Ribose) Links the DNA Damage Response and Biomineralization. Cell Rep 27, 3124–3138 e3113 (2019).

67 Salsas-Escat, R., Nerenberg, P. S. & Stultz, C. M. Cleavage site specificity and conformational selection in type I collagen degradation. Biochemistry 49, 4147–4158 (2010).

68 Hu, J. et al. Design of synthetic collagens that assemble into supramolecular banded fibers as a functional biomaterial testbed. Nat Commun 13, 6761 (2022).

69 den Dunnen, J. T. & Antonarakis, S. E. Mutation nomenclature extensions and suggestions to describe complex mutations: a discussion. Hum Mutat 15, 7–12 (2000).

70 den Dunnen, J. T., et al. HGVS Recommendations for the Description of Sequence Variants: 2016 Update. Hum Mutat 37, 564–569 (2016).

71 Carafoli, F., Hamaia, S. W., Bihan, D., Hohenester, E. & Farndale, R. W. An activating mutation reveals a second binding mode of the integrin alpha2 I domain to the GFOGER motif in collagens. PLoS One 8, e69833 (2013).

72 Orgel, J. P., Irving, T. C., Miller, A. & Wess, T. J. Microfibrillar structure of type I collagen in situ. Proc Natl Acad Sci U S A 103, 9001–9005 (2006).

