## Supplementary Information for "Nanoscale Structural and Functional Impacts of Disease-Associated Collagen Mutations"

### Supplementary Figures

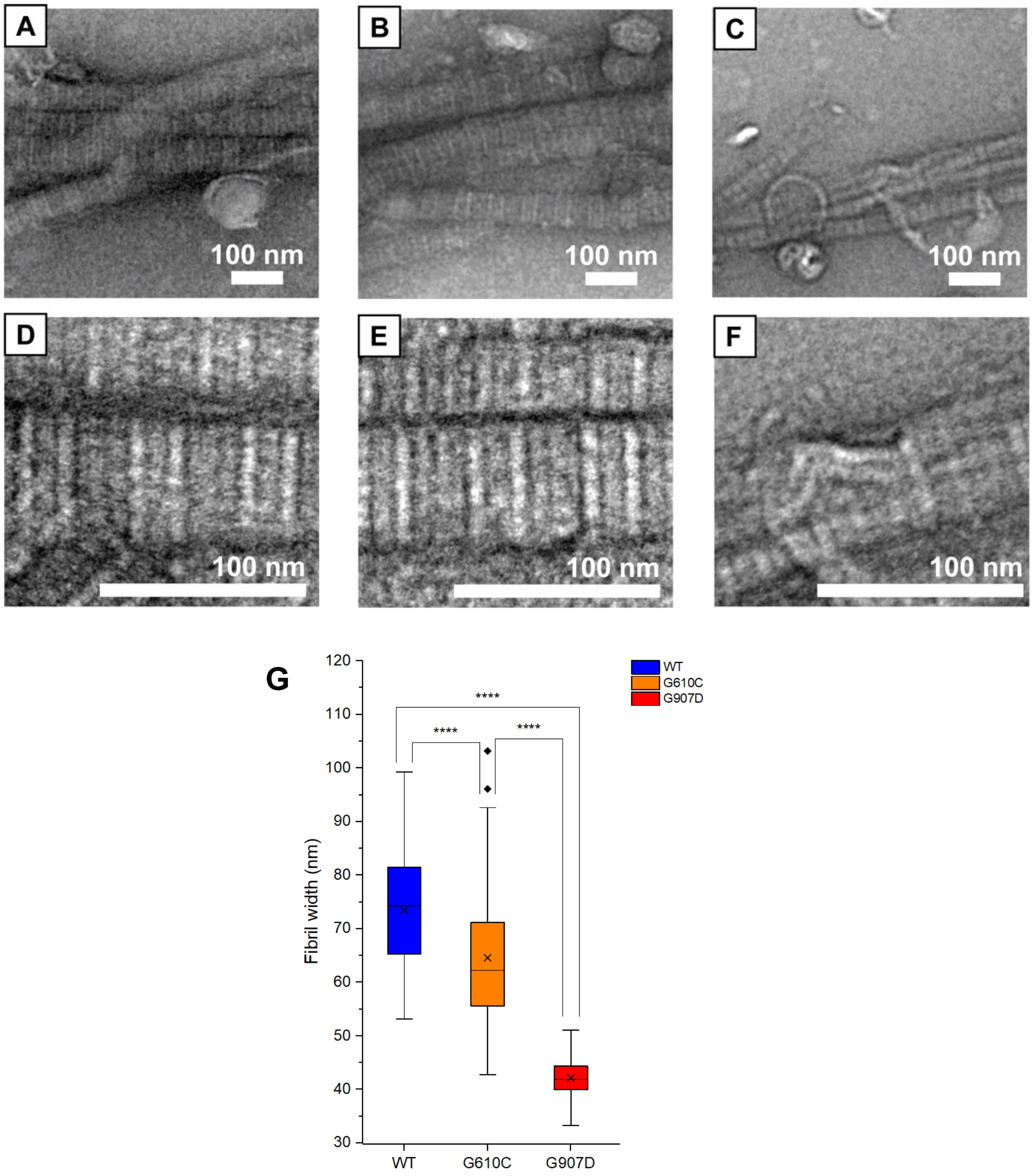


**Figure S1: BF-TEM images of negatively stained collagen fibrils from ECM produced by human fibroblasts.** BF-TEM images from negatively stained (A,D) WT, (B,E) G610C, and (C,F) G907D collagen fibrils. All scalebars = 100nm. (A-C) Fibrils at 8000x magnification. (D-F) Fibrils at 30,000x magnification. (G) Statistical analysis of the width of the fibrils.

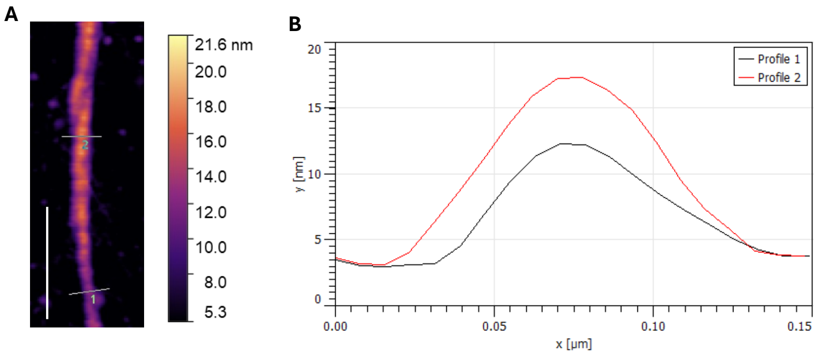


**Figure S2: Local morphological changes in G907D fibrils.** (A) AFM image of a G907D fibril (scale bar: 500 nm) showing the locations from which cross-sectional height profiles were extracted. (B) Cross-sectional profiles reveal variations in height and width along the same fibril, indicating localized fibril thinning. Profile 1 (black) corresponds to a narrower region of the fibril, whereas Profile 2 (red) corresponds to a broader region, confirming localized thinning.

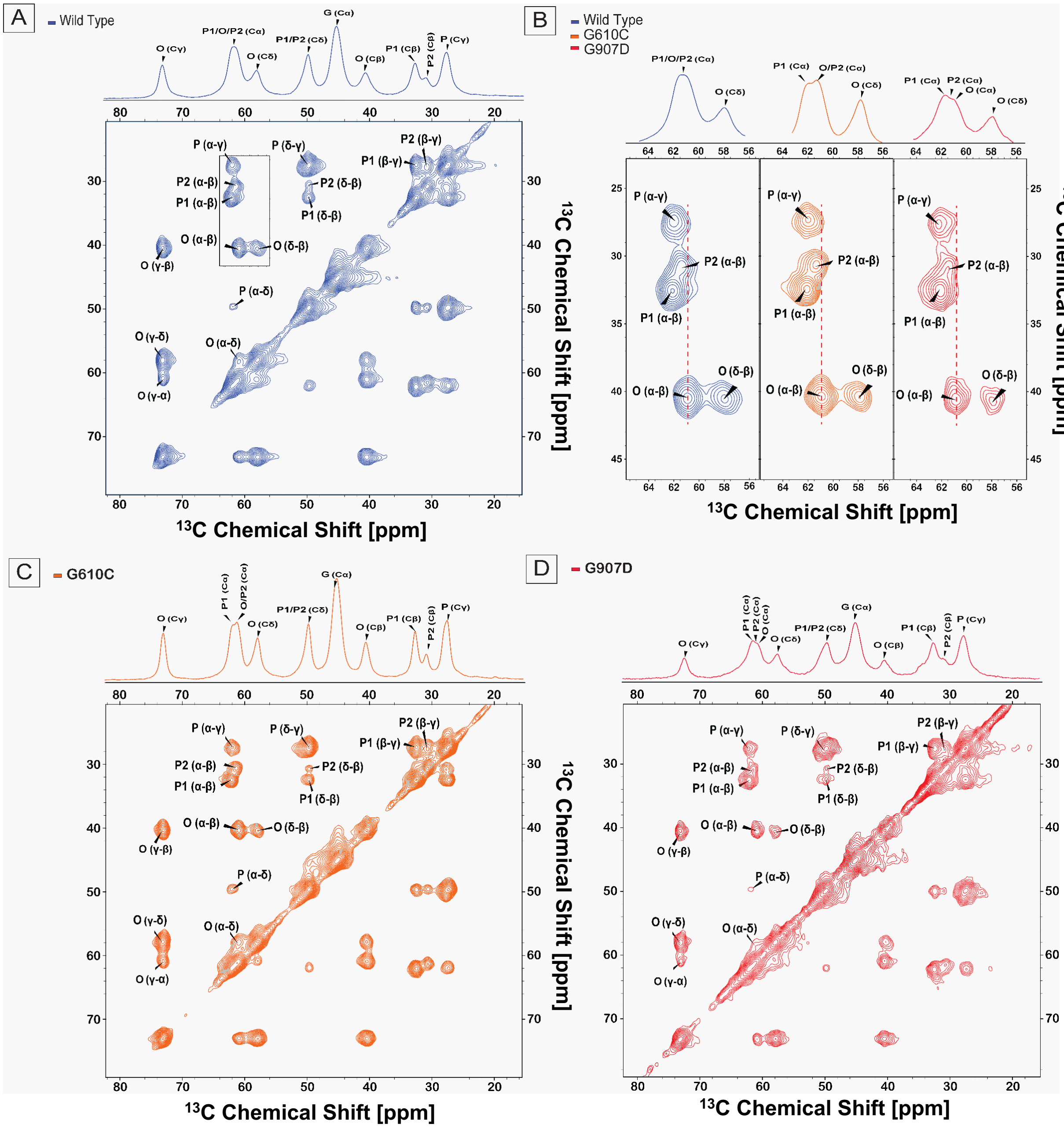


**Figure S3: 2D $^{13}C$-$^{13}C$ DARR Spectra of human fibroblast ECM collagen fibrils at 25°C.** (**A**) DARR spectra for Wild Type (blue), (**B**) An enlarged view of the Pro/Hyp Cα cross-peaks for the Wild Type (blue), G610C (orange), and G907D (red) is shown in the slice, (**C**) DARR spectra for G610C (orange), (**D**) DARR spectra for G907D (red). 1D $^{1}H$-$^{13}C$ CP spectra are shown at the top of the figures. The contour levels for 2D experiments were scaled for better visual comparison and reflected an accurate comparison of the intensities between experiments.

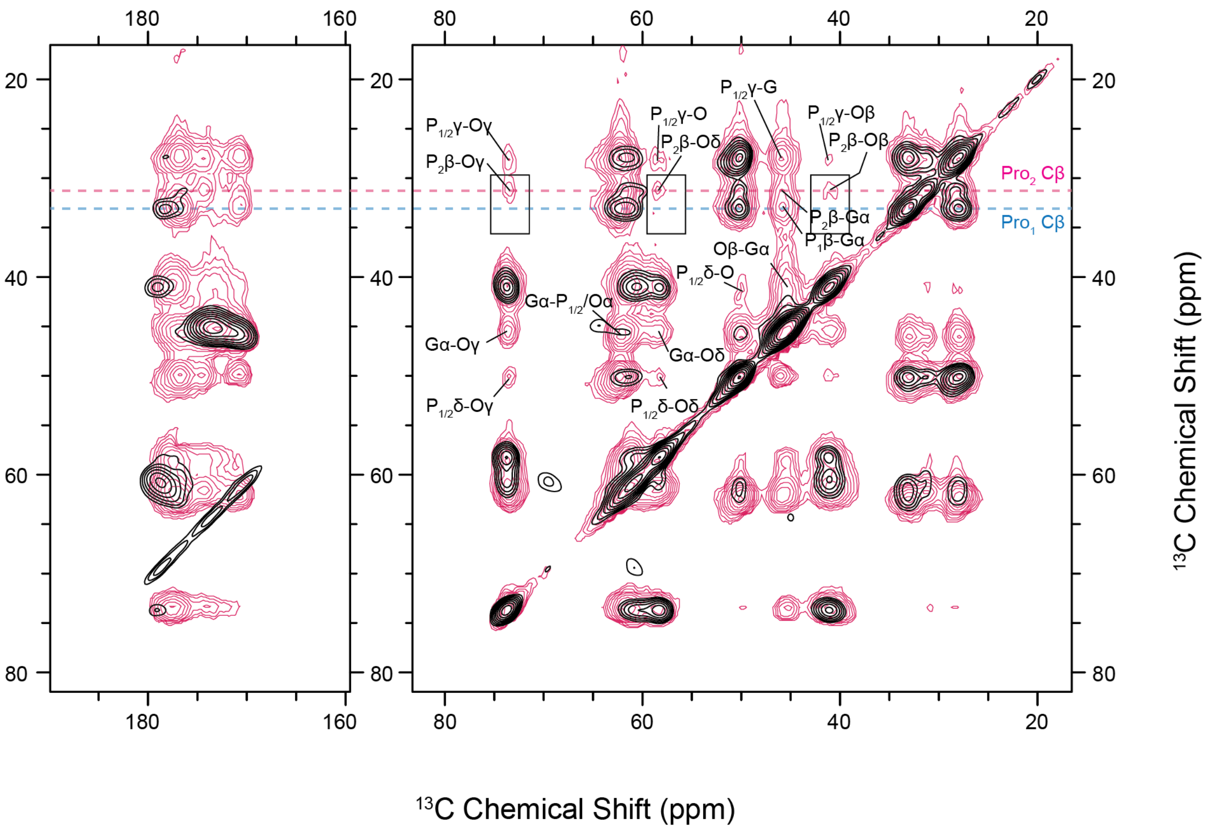


**Figure S4: Assignment of $P_{GPO}$ and $P_{GPX}$.** 2D $^{13}C$-$^{13}C$ DARR spectra of [U-$^{13}C$,$^{15}N$]-Gly, Pro, Hyp-ECM from MC3T3 cells. $^{13}C$-$^{13}C$ DARR experiments acquired with $^{13}C$-$^{13}C$ mixing times of 100 ms (pink), in which inter-residue crosspeaks are visible, and 8 ms (black), in which only intra-residue crosspeaks are detected. In the case of $Pro_2$ (pink dashed line), crosspeaks between $P_2C\beta$—$OC\beta$, $P_2C\beta$—$OC\gamma$, and $P_2C\beta$—$OC\delta$ are observed with 100 ms $^{13}C$-$^{13}C$ mixing (boxed). However, these inter-residue crosspeaks are not observed for $Pro_1C\beta$ (blue dashed line). Therefore, $Pro_2$ is attributed to Pro residues preceding Hyp residues ($P_{GPO}$), whereas $Pro_1$ is attributed to Pro residues not preceding Hyp residues ($P_{GPX}$).

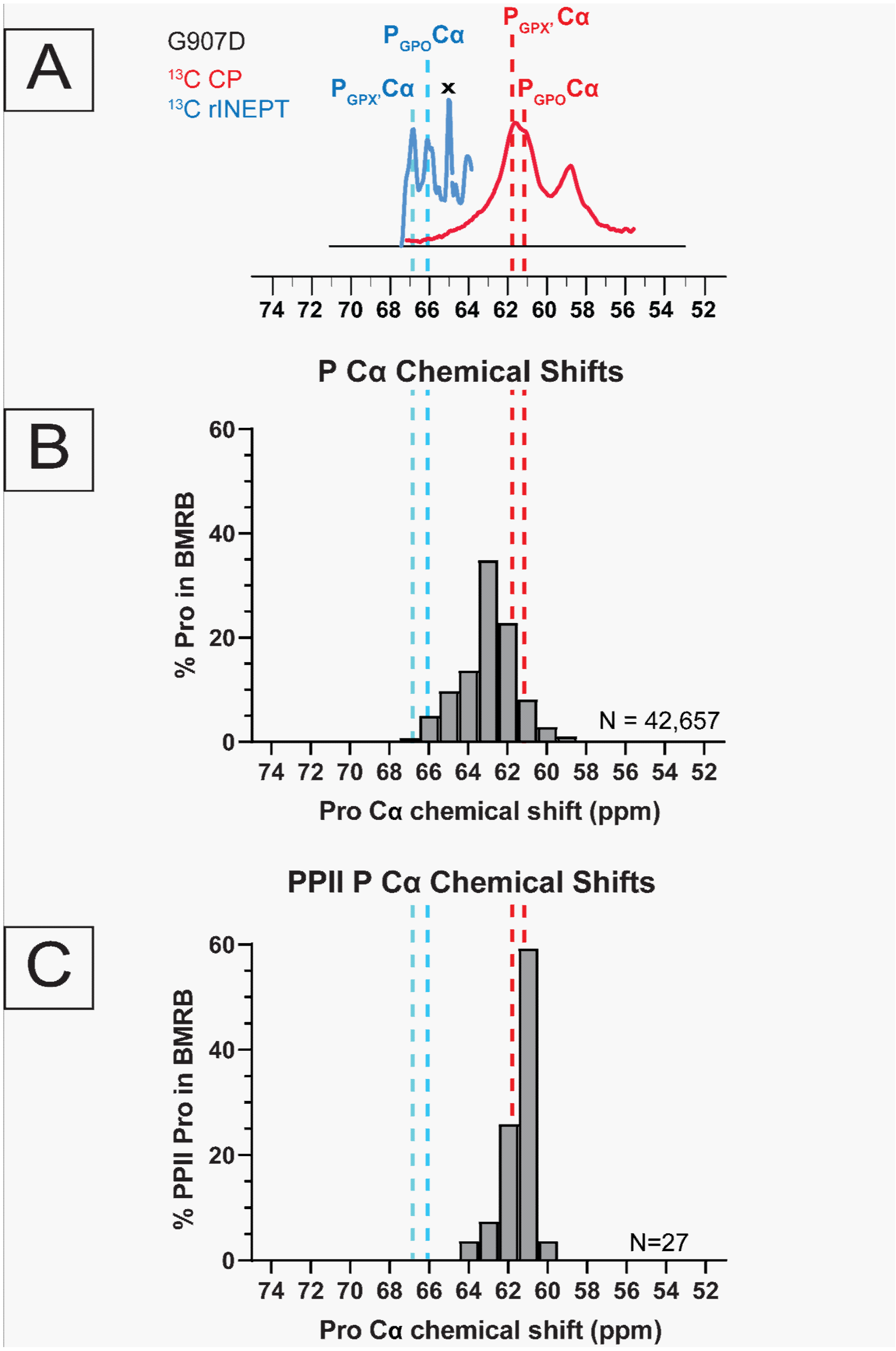


**Figure S5: Determination of ECM Pro secondary structure based on Cα chemical shifts.** (A) Cα regions of 1D $^{1}H$-$^{13}C$ CP (red) and $^{13}C$ INEPT (cyan) spectra overlayed. Experiments were performed at 25°C. The $^{13}C$-INEPT spectrum is amplified 3x for clarity. The dotted lines draw the eye from the assigned ECM Pro Cα peaks to their positions in the histograms from the BMRB. (B-C) Histograms show populations of the Pro Cα chemical shifts found in the BMRB. Each bar represents chemical shifts from X.0000 to X.9999. (B) Populations of Pro Cα chemical shifts found within all BMRB entries. Typical Pro Cα chemical shifts range from 59-67 ppm. (C) Populations of Cα chemical shifts of Pro within PPII secondary structures. The typical range of these Cα chemical shifts is between 60-64 ppm. The Pro Cα chemical shifts in the $^{1}H$-$^{13}C$ CP-based spectra align well with the highest population of PPII-like Pro Cα chemical shifts (red dashed line). This is consistent with a high population of PPII-like Pro in the triple helical regions of the collagen within the ECM. Conversely, the Pro Cα chemical shifts from the $^{13}C$-INEPT-based spectra have very unique downfield chemical shifts, suggesting that these distinctly mobile Pro bear a unique structure.

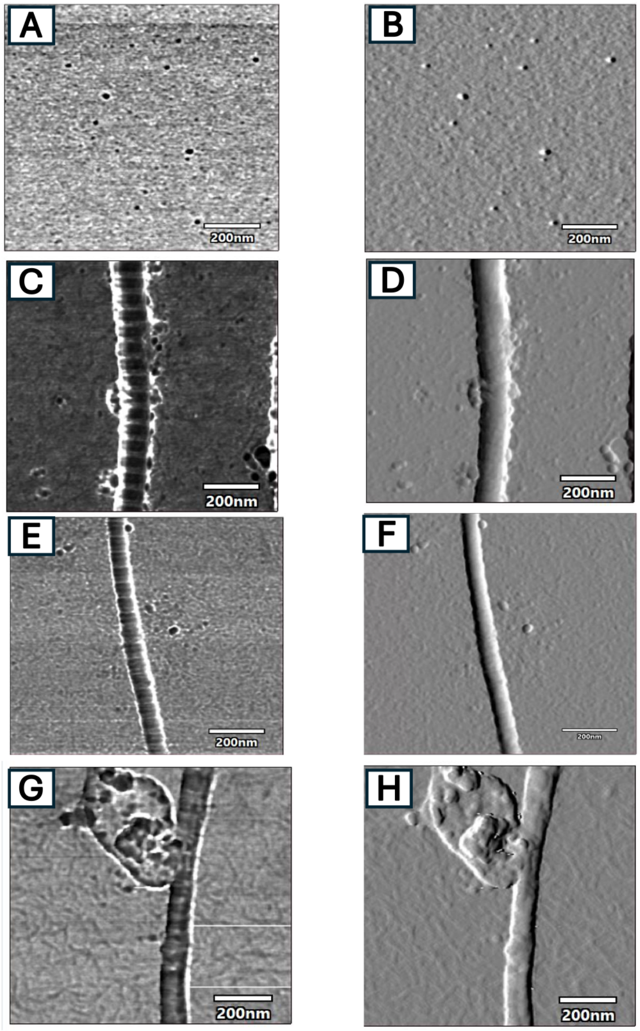


**Figure S6: AFM images for collagen- $\alpha_2$I integrin complexes.** AFM phase (first column) and amplitude (second column) images of collagen- $\alpha_2$I integrin complexes. (A-B) $\alpha_2$I integrin only, (C-D) WT collagen - $\alpha_2$I integrin, (E-F) G610C collagen - $\alpha_2$I integrin and (G-H) G907D collagen - $\alpha_2$I integrin.

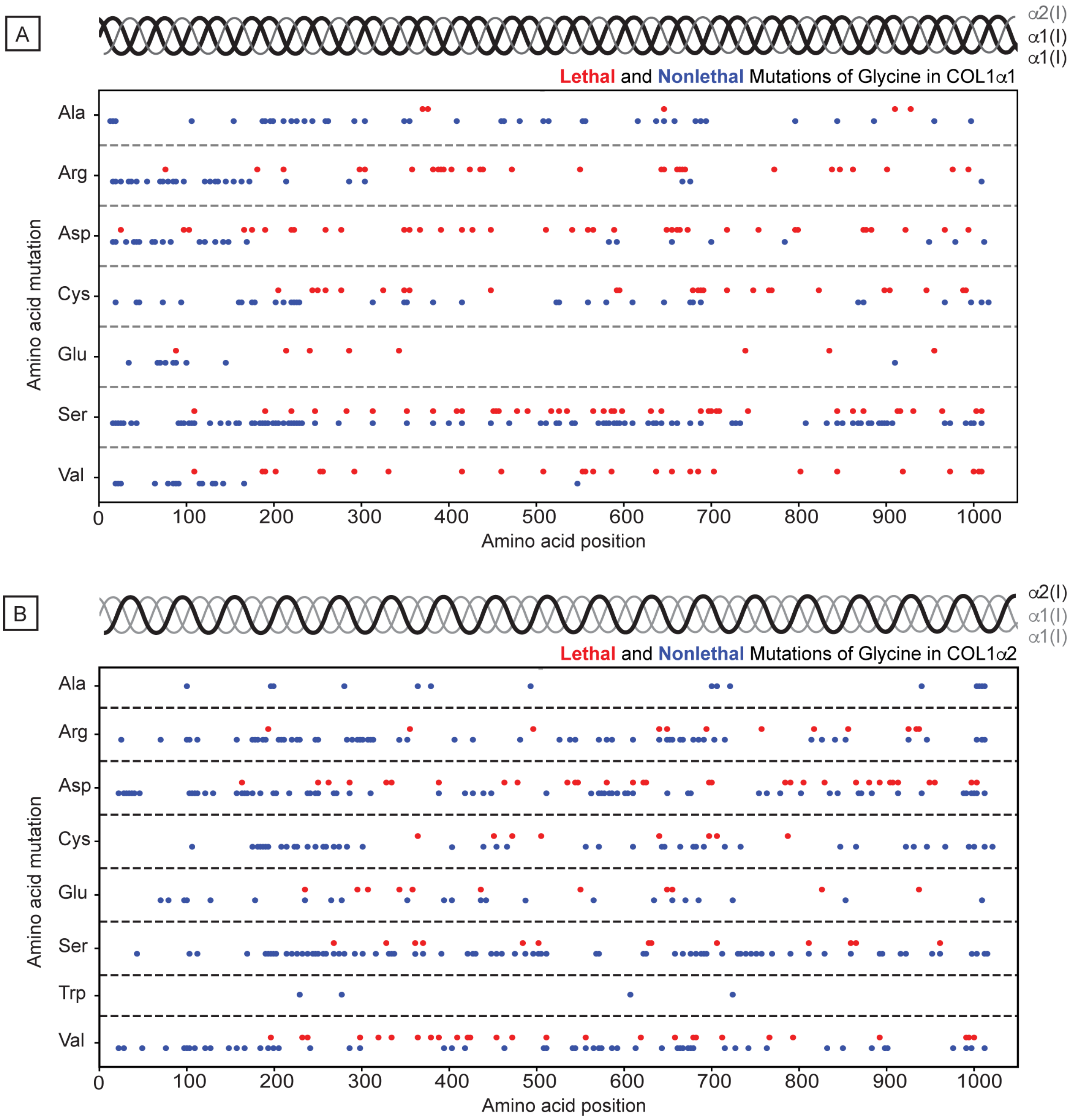


**Figure S7: Distribution of OI Gly missense mutations along the collagen α1(I) and α2(I) sequences.** Analysis of the collagen sequence according to location of OI mutations along the collagen I chains, excluding telopeptides and propeptides. Dots are colored red and blue for lethal and non-lethal mutations, respectively. **(A)** Dot plot of different amino acids showing the distribution of mutations along the collagen I α1 chains. **(B)** Dot plot of different amino acids showing the distribution of mutations along the collagen I α2 chain.

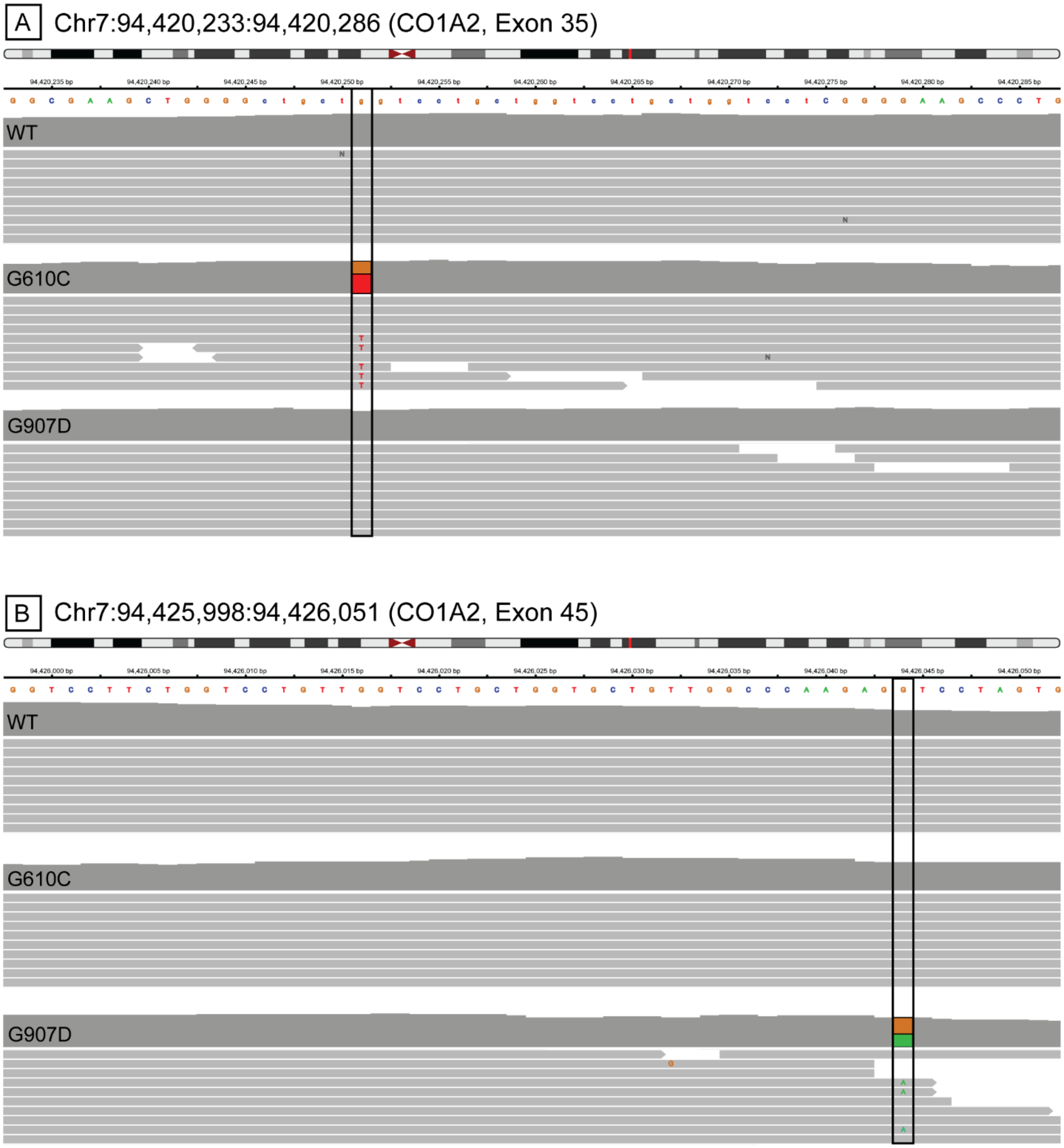


**Figure S8. Alignment results from NGS Whole Exome sequencing shows that the mutant fibroblast cells contained the correct respective mutations.** WT, G610C, and G907D cell lines were sequenced and analyzed to determine the presence of mutations in the collagen chains. The sequence alignments shown are for the collagen α2 chain on chromosome 7. Not all alignments are shown, since most contain over 40 alignment sequences per sample. (A) Position 94,420,233-94,420,286 corresponds to the locations of exons 35 which is expected to contain the G610C OI mutations. In the “G610C” cell line, position 94,420,251 contains the G>T nucleotide mutation which results in Gly to Cys mutations. 60% of the total sequence was mutated to thymine, showing that it contained the correct mutation in a single CO1A2 allele. The “WT” and “G907D” cell lines, as expected, do not contain the G610C mutation. (B) Position 94,425,998-94,426,051 corresponds to the location of exon 45 which is the site which would contain the G907D mutation. G907D mutations are present in only the “G907D” cell lines at position 94,426,044, containing a G>A nucleotide mutation, resulting in a Gly to Asp mutation. In the “G907D” cell line, 45% of the sequences were mutated to adenine, showing the G907D line contains both the correct mutation and is present in only a single CO1A2 allele, as anticipated. The G907D mutation is not present in the “WT’ or “G610C” lines.

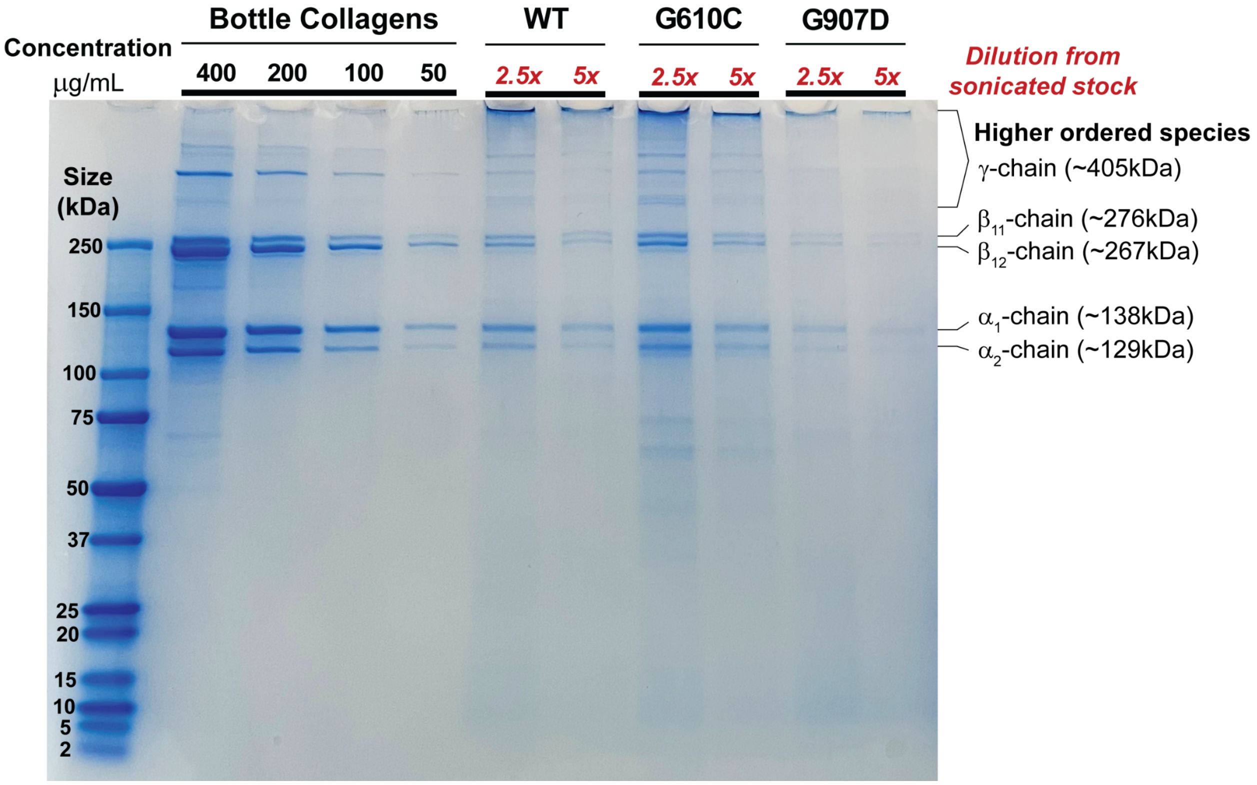


**Figure S9. SDS-PAGE Gel showing concentrations of fibrillar collagens relative to bottled collagens.** WT, G610C, and G907D collagen fibrils were diluted from the sonicated stock solutions and run on an SDS-PAGE Gel with rat tail tendon bottled collagen to determine approximate concentrations of the collagen fibrils. The ladder used was able to determine bands up to approximately 250kDa which includes the collagen α-chains and β-chains (containing one each of α1 and α2 collagen chains, or two α1 chains). Higher order collagen species, including the γ-chain (triple helical collagen) were also shown but not specifically labeled. The lanes which contain denatured collagen fibrils all contain proteins which were unable to travel through the gel and thus, a portion of fibrils present cannot be accounted for in the final concentrations. The final concentrations derived from the concentrations shown in the gel were 250 μg/mL WT collagen, 375 μg/mL G610C collagen, and 125 μg/mL G907D collagen.

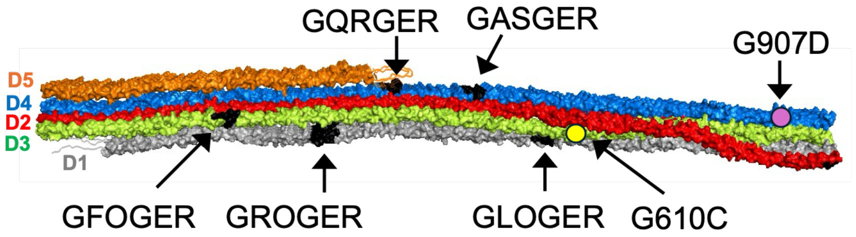


**Figure S10. Spatial location of integrin binding sites on a D-period.** Additional binding sites other than GFOGER and relative location of G610C and G907D mutation.

**Table S1: Summary of statistics from AFM fibril analyses.** Mean heights and periodicities of WT, G610C, and G907D fibrils were obtained from AFM images. Statistical significances were analyzed via GraphPad/PRISM software. The N (number of fibrils analyzed), mean, and standard deviation of each of the datasets are summarized in the table. “ns” denotes non-significant.

| | Periodicity | | Heights | |
|---|---|---|---|---|
| | Data Significance | P-value | Data Significance | P-value |
| WT to G610C | ns | 0.9865 | ns | 0.2514 |
| WT to G907D | ns | 0.9922 | Yes **** | <0.0001 |
| G610C to G907D | ns | 0.9704 | Yes *** | 0.0002 |

| | N | Mean periodicity (nm) | N | Mean Heights (nm) |
|---|---|---|---|---|
| WT | 15 | 65.2 ± 0.7 | 19 | 19.5 ± 4.3 |
| G610C | 21 | 65.3 ± 0.8 | 25 | 17.3 ± 4.1 |
| G907D | 15 | 65.1 ± 1.9 | 18 | 12.0 ± 3.5 |

**Table S2: Summary of statistical analysis of TEM images of fibrils.** Mean widths of WT, G610C and G907D fibrils were obtained from TEM images. The N (number of measurements), mean, and standard deviation of each of the datasets are summarized in the table. "ns" denotes non-significant.

| | Width | |
|---|---|---|
| | N | Mean Width (nm) |
| WT | 100 | 73.5 ± 1.0 |
| G610C | 97 | 65.6 ± 1.3 |
| G907D | 25 | 42.2 ± 0.9 |

| | Data Significance | P-value |
|---|---|---|
| WT to G610C | Yes **** | <0.0001 |
| WT to G907D | Yes **** | <0.0001 |
| G610C to G907D | Yes **** | <0.0001 |

**Table S3. $^{13}$C Chemical shifts of ECM variants derived from CP-based or INEPT-based experiments.** Reported chemical shifts are taken from 1D $^{1}$H-$^{13}$C and 2D $^{13}$-$^{13}$C DARR experiments (CP-based) and 1D $^{13}$C and 2D $^{13}$C-$^{13}$C TOBSY experiments (INEPT-based). $^{13}$C chemical shifts are referenced to DSS.

| | **CP-based** | | | **INEPT-based** |
|---|---|---|---|---|
| Assignment | WT | G610C | G907D | G907D |
| G1 Cα | 45.07 | 45.08 | 45.12 | 45.59 |
| G1 CO | 170.26 | 170.30 | 170.34 | -- |
| G2 Cα | 45.07 | 45.08 | 45.12 | -- |
| G2 CO | 172.73 | 172.8 | 172.78 | -- |
| P1 Cα | 61.50 | 61.76 | 61.88 | 66.58 |
| P1 Cβ | 32.44 | 32.41 | 32.38 | 32.42 |
| P1 Cγ | 27.41 | 27.29 | 27.46 | 27.46 |
| P1 Cδ | 49.62 | 49.61 | 49.83 | 50.49 |
| P1 CO | 174.10 | 174.06 | 174.03 | -- |
| P2 Cα | 61.50 | 61.23 | 61.18 | 65.64 |
| P2 Cβ | 30.67 | 30.61 | 30.78 | 30.79 |
| P2 Cγ | 27.41 | 27.29 | 27.46 | 27.46 |
| P2 Cδ | 49.62 | 49.61 | 49.83 | 50.49 |
| P2 CO | 176.30 | 176.30 | 176.40 | -- |
| O Cα | 61.50 | 61.23 | 61.40 | 65.83 |
| O Cβ | 40.41 | 40.35 | 40.41 | 40.29 |
| O Cγ | 73.01 | 73.02 | 73.01 | 73.41 |
| O Cδ | 57.86 | 57.82 | 57.76 | 58.11 |

| | | | | |
|---|---|---|---|---|
| O CO | 177.30 | 177.40 | 177.20 | -- |

**Table S4. Summary of statistical analysis of AFM data on integrin binding to collagen fibrils.** Contour length of the collagen fibrils were measured with ImageJ software, while the integrin numbers were determined from the globular features bound to collagen fibrils. Phase and amplitude AFM images were considered for identification of integrin contours on collagen fibrils.

| Sample | Number of collagen fibrils analyzed | Total contour length of collagen fibril analyzed(nm) | Number of integrins per micron collagen |
|---|---|---|---|
| WT-ECM | 10 | 10371.6 | 5.49±1.3 |
| G610C | 16 | 16058.8 | 2.36±0.5 |
| G907D | 10 | 10036.6 | 9.76±0.9 |
| WT-ECM without $Mg^{2+}$ | 10 | 12121.0 | 1.1±0.4 |

| | Data significance for number of integrins per micron collagen | p-value |
|---|---|---|
| WT-ECM to G610C | Yes | 0.02 |
| WT-ECM to G907D | Yes | 0.017 |
| WT-ECM to WT-ECM without $Mg^{2+}$ | Yes | 0.004 |

**Table S5.** Odds ratio analysis comparing lethal (Type II) and nonlethal (Types I, III, IV) cases of Osteogenesis Imperfecta for different Glycine mutations on collagen I genes by resultant amino acid and alpha 1 or alpha 2 chain. Most of the results are significant and illustrate that alpha 1 mutations are generally more lethal. Statistics are calculated based on the null hypothesis that an Odds Ratio is equal to one.

| | Lethal A1 Cases | Nonlethal A1 Cases | Lethal A2 Cases | Nonlethal A2 Cases | Odds Ratios (OR) | Confidence Intervals (OR) | lnOR | Standard Error (lnOR) | P Values |
|---|---|---|---|---|---|---|---|---|---|
| **Ala** | 6 | 54 | 0 | 22 | 5.37 | 0.29 - 99.3 | 1.68 | 1.49 | 2.59e-01 |
| **Arg** | 58 | 101 | 12 | 102 | 4.88 | 2.47 - 9.63 | 1.59 | 0.35 | 4.85e-06 |
| **Asp** | 42 | 40 | 39 | 124 | 3.34 | 1.90 - 5.86 | 1.21 | 0.29 | 2.71e-05 |
| **Cys** | 37 | 88 | 10 | 69 | 2.90 | 1.35 - 6.24 | 1.07 | 0.39 | 6.45e-03 |
| **Glu** | 10 | 13 | 14 | 37 | 2.03 | 0.73 - 5.69 | 0.709 | 0.52 | 1.76e-01 |
| **Ser** | 67 | 393 | 28 | 388 | 2.36 | 1.49 - 3.75 | 0.860 | 0.24 | 2.72e-04 |
| **Val** | 38 | 40 | 32 | 90 | 2.67 | 1.47 - 4.87 | 0.983 | 0.31 | 1.32e-03 |
| **Total** | 258 | 729 | 135 | 832 | 2.18 | 1.73 - 2.75 | 0.780 | 0.12 | 3.48e-11 |

**Table S6.** Compilation of significant results from odds ratio analysis comparing lethal (Type II) and nonlethal (Types I, III, IV) cases of Osteogenesis Imperfecta for different Glycine mutations on collagen I genes by resultant amino acid (irrespective of alpha chain). Statistics are calculated with a null hypothesis that an Odds Ratio is equal to one.

| | | Lethal Cases | Nonlethal Cases | Odds Ratios (OR) | Confidence Intervals (OR) | lnOR | Standard Error (lnOR) | P Values |
|---|---|---|---|---|---|---|---|---|
| **Arg** | Arg | 70 | 203 | 4.37 | 1.82 – 10.5 | 1.47 | 0.446 | 9.51e-04 |
| | Ala | 6 | 76 | | | | | |
| | Arg | 70 | 203 | 2.83 | 2.01 – 4.00 | 1.04 | 0.176 | 3.29e-09 |
| | Ser | 95 | 781 | | | | | |
| **Asp** | Asp | 81 | 164 | 6.26 | 2.61 – 15.0 | 1.83 | 0.445 | 3.82e-05 |
| | Ala | 6 | 76 | | | | | |
| | Asp | 81 | 164 | 1.65 | 1.08 – 2.51 | 0.501 | 0.215 | 1.97e-02 |
| | Cys | 47 | 157 | | | | | |
| | Asp | 81 | 164 | 4.06 | 2.89 – 5.71 | 1.40 | 0.174 | 8.88e-16 |
| | Ser | 95 | 781 | | | | | |
| **Cys** | Cys | 47 | 157 | 3.79 | 1.55 – 9.26 | 1.33 | 0.455 | 3.43e-03 |
| | Ala | 6 | 76 | | | | | |
| | Cys | 47 | 157 | 2.46 | 1.67 – 3.63 | 0.901 | 0.199 | 5.78e-06 |
| | Ser | 95 | 781 | | | | | |
| **Glu** | Glu | 24 | 50 | 6.08 | 2.32 – 15.9 | 1.81 | 0.491 | 2.40e-04 |
| | Ala | 6 | 76 | | | | | |
| | Glu | 24 | 50 | 3.95 | 2.32 – 6.71 | 1.37 | 0.271 | 4.10e-07 |
| | Ser | 95 | 781 | | | | | |
| **Val** | Val | 70 | 130 | 6.82 | 2.83 – 16.5 | 1.92 | 0.449 | 1.92e-05 |
| | Ala | 6 | 76 | | | | | |
| | Val | 70 | 130 | 1.56 | 1.05 – 2.32 | 0.446 | 0.203 | 2.81e-02 |
| | Arg | 70 | 203 | | | | | |
| | Val | 70 | 130 | 1.80 | 1.16 – 2.78 | 0.587 | 0.223 | 8.41e-03 |
| | Cys | 47 | 157 | | | | | |
| | Val | 70 | 130 | 4.43 | 3.09 – 6.35 | 1.49 | 0.184 | 6.66e-16 |
| | Ser | 95 | 781 | | | | | |

### Supplementary Method

#### OI Mutation Database Analyses

Beyond basic visualizations of the cleaned mutation data that qualitatively represent the effect of mutation location, odds ratio analyses provide quantitative evidence of statistically significant differences between outcomes for two types of mutations (the odds of something occurring in one group and the odds of it occurring in a different group[1]. For the collagen I mutation data, these odds ratio analyses compare the number of lethal and nonlethal cases between two different groups such as alpha I mutations and alpha II mutations or Gly to Ser and Gly to Val mutations. The following equations exhibit our process of calculating odds ratios and relevant statistical values.

***Odds Ratio (OR)***

| | Lethal Cases | Nonlethal Cases |
|---|---|---|
| Group 1 | $a$ | $b$ |
| Group 2 | $c$ | $d$ |

$$Odds\ of\ Group\ 1 = \frac{a}{b}$$

$$Odds\ of\ Group\ 2 = \frac{c}{d}$$

***Odds Ratio (OR) (cont.)***

$$OR = \frac{\left(\frac{a}{b}\right)}{\left(\frac{c}{d}\right)}$$

***Log Odds Ratio (ln(OR))***

$$\ln(OR) = \ln\left(\frac{\left(\frac{a}{b}\right)}{\left(\frac{c}{d}\right)}\right)$$

***Standard Error of Log Odds Ratio (SE ln(OR))***

$$SE\ \ln(OR) = \sqrt{\left(\frac{1}{a}\right) + \left(\frac{1}{b}\right) + \left(\frac{1}{c}\right) + \left(\frac{1}{d}\right)}$$

***Odds Ratio Confidence Intervals (OR CIs)***

$$OR\ CIs\ =\ e^{(ln(OR)\pm\ 1.96\times(SE\ ln(OR))}$$

***Z-Score***

$$z = \frac{ln(OR)}{SE\ ln(OR)}$$

***P-Value***

$$p =\ 2 \times \left(1 - P(Z \geq |z|)\right)$$

For odds ratios, statistical values like standard error or 95 percent confidence intervals are calculated and interpreted slightly differently than more traditional cases, given that the null hypothesis for an odds ratio analysis is that the odds ratio is equal to one–meaning that the odds for one group are the same as the odds for the second group[2]. In scenarios where no cases are present (e.g. no lethal type II OI cases are present for a particular group) and mathematical operations using zero become a problem while calculating odds ratios, adding 0.5 to the a, b, c, and d values representing cases of each group is the standard remedy. Odds ratios were always calculated such that the larger odds were divided by the smaller odds for consistency and to support the intuition of making statements to the effect that one group is however many times more lethal than another.

**References:**


1 Bond, S. & FU, B. Essential Medical Statistics (2nd edn). Kirkwood BR, Sternc JAC. Malden, MA: Blackwell Publishing, 2003, pp. 288, $52.95 (PB). ISBN 0865428719. *International Journal of Epidemiology* **33**, 1418-1419 (2004).

2 Ashby, D. Practical statistics for medical research. Douglas G. Altman, Chapman and Hall, London, 1991. No. of pages: 611. Price: £32.00. *Statistics in Medicine* **10**, 1635-1636 (1991).
